# DNA cross-over motifs based, programmable supramolecular hydrogels for mechanoregulatory effects of cellular behaviour and cytoskeleton reorganization

**DOI:** 10.1101/2025.05.14.653944

**Authors:** Ankur Singh, Dhiraj Bhatia

## Abstract

The development of mechanically tunable DNA-based supramolecular hydrogels that recapitulate the dynamic mechanoelastic properties of native extracellular matrices (ECMs) is critical for advancing 3D tissue engineering. DNA supramolecular hydrogels with their precise programmable architecture and impeccable control at the nanometre scale assembly make them a highly lucrative biopolymer in the design of ECM-mimicking scaffolds. This study explores the fabrication of stiffness-customizable rigid DNA hydrogels with varying branching architectures, including Double Crossovers (DX), Paranemic Crossovers (PX), and Tensegrity motif structures with three to six arms. Our study reveals that by incorporating different numbers of crossovers, branching arms, and the type of sticky-armed linkers, it is possible to generate a range of DNA hydrogels with varying network geometry and mechanoelastic properties ranging from 50-300 kPa, without any chemical / enzymatic crosslinking aid. These DNA hydrogels mechanoregulate cellular behaviour in retinal pigment epithelial (RPE1) cells by enhancing cellular adhesion, elongation (3 - 8x area increase vs. control), viability (concentration-dependent, 1 - 8x vs. control), organellar homeostasis, including mitochondrial fragmentation and ER stress attenuation. This work establishes a framework for automating DNA hydrogel-based scaffold fabrication processes to meet the preferences of various cells/tissues with bespoke mechanical cues, advancing personalized high-throughput tissue engineering platforms.

**Graphical abstract:** 

## I. Introduction

Hydrogels have already proven their indispensable role in tissue engineering by providing a three-dimensional, porous, and hydrated ECM-mimicking microenvironment for the comprehensive growth and development of cells and tissues(Zhu and Marchant, 2011). Moreover, conventional hydrogels like collagen and alginate lack the mechanical tunability required to match the mechanoelastic needs of diverse human tissues (e.g., brain and adipose tissues: ∼0.1-1 kPa; cartilage: ∼10-25 kPa; skin: ∼10-100 kPa; bone: >1 MPa)(Guimarães *et al*., 2020; Conway *et al*., 2023). This limitation impedes the use of conventional hydrogels in regulating cell fates, influencing their adhesion, differentiation, and organellar functions in response to the mechanoelasticity of the scaffolds(Engler *et al*., 2006; Walters and Gentleman, 2015).

DNA hydrogels offer unique advantages, both in form and function, by their inherent biocompatibility, biodegradability, along with biological functionality coupled with atomic-level precision through sequence design(Seeman, 1982; Wang *et al*., 1991; Mao *et al*., 2000; Yan *et al*., 2002; Aldaye *et al*., 2010). Recent advancements have demonstrated their efficiency and potential in providing ECM-mimicking environments for tissue engineering. Many research groups have demonstrated diverse ways of forming DNA-based hydrogels with mechanical stiffness in the range of 100 Pa - 100 kPa (Luo *et al*., 2022; Lachance-Brais *et al*., 2023; Shi *et al*., 2024; Singh *et al*., 2024; Wang *et al*., 2024). The growing interest in DNA hydrogels stems from their inherent biological properties and ease of customization, offering a platform for modulation of cellular responses in ways that conventional hydrogels cannot.

The architecture of DNA hydrogels is fundamentally determined by the interconnectivity of constituent DNA junctions, which regulate the branching architecture and stiffness of the hydrogels. At the molecular level, these junctions serve as a structural node that defines the network topology. The properties of hydrogel can be programmed by regulating these junctions. In our previous work, we tried to explore the role of soft DNA hydrogels made from branched simple DNA duplex junctions in programming the cellular behaviour of RPE1 cells, including enhanced cellular spreading, elongation, endocytic uptake, perturbation of ECM-regulating surface markers, mitochondrial fragmentation, and focal adhesion complex regulation (Singh *et al*., 2024; Singh, Singh and Bhatia, 2024). These hydrogels showed the mechanical range of 0.1 Pa – 10 Pa, without any addition of crosslinkers or chemical modifications. Extending the scope of our previous work, reinforcing simple duplex junctions with mechanically robust and topologically stable junctions may serve as a key to unlock more features for DNA-based hydrogels in designing scaffolds for automated 3D bioprinting while systematically probing mechanobiology.

While prior studies have explored DNA hydrogels for drug-delivery or biosensing, their potential as a mechanically tunable scaffold for tissue engineering applications remains underexplored. Integration of more complex DNA motif structures such as i-motifs, G-quadruplexes, DNA crossovers, etc., into junctions may lead to diversification of geometry and mechanical properties. DNA motifs such as double-crossovers (DX) contain two crossover sites between helical domains, while a paranemic-crossover (PX) features an even more complex design, featuring a four-stranded coaxial DNA complex with crossovers happening at every possible point (Rodriguez *et al*., 2025). These structures must be designed while keeping the topology of DNA strands in mind, where crossovers are separated by specific half-turns to avoid torsional stress. The increased structural rigidity of DX, and even so for PX, compared to simple duplexes, makes them valuable building blocks for constructing more stable DNA nanostructures and hydrogels (Rodriguez *et al*., 2025; Scarton *et al*., 2025). Increasing the degree of sustained crossovers is one way to increase the rigidity of the hydrogels.

Another significant advancement is the development of hydrogels with tensegrity architectures inspired by the mechanochemical principles underlying cell-ECM interactions(Lu *et al*., 2021, 2023; Xue *et al*., 2025). These motifs are made from ssDNA strands forming an isolated rigid compression component, held in place by continuous network of tensile elements, enabling tensile moduli up to 30 MPa (Xue *et al*., 2025). The incorporation of Tensegrity principles into DNA hydrogel design represents a biomimetic approach to create a robust, adaptive platform for advanced mechanoregulated tissue engineering.

We hypothesize that branching complexity via regulating the degree of crossovers and the branching arms of tensegrity structures (DX<PX< Tensegrity (trimers-hexamers)), directly governs hydrogel stiffness, which in turn can modulate cellular mechanotransduction pathways. Hereby, in this work, we have demonstrated design, synthesis, characterization, and applications of DNA hydrogels using DX, PX, and tensegrity motifs (3-6 arms) in programming RPE1 cells’ morphology, viability, and organellar dynamics in a mechanotransduction-dependent manner. Our results highlight the importance of branching units in regulating the stiffness of the hydrogel scaffolds. The truncated DX, PX, and tensegrity motifs (intrinsic controls) showed no potential for forming scaffolds. We also demonstrated a range of fully customizable rigid DNA hydrogel systems, with varying stiffness within the range of 50 kPa - 300 kPa. These hydrogels showed rigidity-dependent cellular programming potentials by enhancing cell area (3-8x compared to poly-L-lysine (PLL)), viability (1-8x compared to PLL), enhanced actin polymerization, and regulation of organellar homeostasis, including mitochondrial fragmentation and Endoplasmic reticulum stress attenuations. Our design strategy enabled the tuneability of DNA hydrogels’ mechanical properties without compromising their structural integrity and biological functionality, thereby highlighting their potential to serve as a customizable platform to guide cellular behaviour while offering a scalable strategy for precision tissue engineering.

## II. Results & Discussions

### a. Design and synthesis of DNA Hydrogels

The Design process of these DNA hydrogels is based on the Watson-Crick base pairing rule. Here we exploit the specific nature of nucleotide base-pairing to our advantage in designing structures with DX, PX, and tensegrity motifs. The sequence design pipeline follows several rules: 1. Each crossover should be separated by integral half-turns of helical coil, to avoid torsional strain. 2. Incorporation of AT-rich sequences near the crossover sites and GC-rich sequences near the terminal ends ensures thermodynamically stable structures. 3. Sequences should be checked to avoid the potential for forming secondary structures or heterodimers. 4. Pair-wise alignment of ssDNA strands must be checked to calculate the minimum Gibbs Free energy (ΔG). 5. Visualization of assembled structures through scadNANO (2D-assemblies) and oxDNA (3D assemblies) must be ensured. 7. Finalized sequences should be immunologically inert (checked via BLAST tools).

We designed fourteen unique DNA nanostructures, eight of which form intricate hydrogel networks, while six act as truncated controls. We also explored the nature of the sticky-arms in regulating the rigidity of the DNA hydrogels. As a result, we have three variants of DX, namely DX0 (truncated motif), DX1 (DX motif with non-palindromic sticky end linkers), and DX2 (DX motif with palindromic linkers). The presence/absence of palindromic linkers affects the scaffold stiffness by regulating the degree of freedom in which sticky-end-based joining is allowed. Palindromic linkers offer higher degrees of freedom compared to the sequence-confined non-palindromic linkage. Similar to DX, there are also three variants of PX, including PX0 (truncated control), PX1 (PX motif - non-palindromic linker type), and PX2 (PX motif - palindromic linker type). However, for tensegrity structures, we decided to only incorporate non-palindromic linkers, thereby giving two variants of Tensegrity triangles (TT0, TT1), Quadrilaterals (TQ0, TQ1), Pentamers (TP0, TP1), and Tensegrity Hexamers (TH0, TH1).

### b. Characterization of DNA hydrogels

The finalized sequences following the above design pipelines were outsourced from Merck. The lyophilized ssDNA strands were dissolved in nuclease-free water (NFW) and stored at 4°C until use. The synthesis of DNA hydrogel was initiated by mixing the ssDNA oligonucleotides in the desired ratio and concentrations (Supplementary Table 1) supplemented with 50 mM of MgCl_2_ as a bridging agent. The assembly of these strands was analysed by performing electrophoretic mobility shift assay (EMSA) on 10% Native PAGE systems.

#### EMSA

The electrophoretic mobility shift assay is a gel-based electrophoresis method where the sample is passed through the tight matrix of a well-defined resolving gel system (e.g., polyacrylamide or agarose) under the influence of an external electric field, thereby providing a sieving effect on the sample. The DNA hydrogel samples were loaded in a systematic manner, where to the left of the DNA ladder lane, ssDNA were loaded, and on the right, the subsequent assembly of these ssDNA strands was shown in a cascading manner. The higher the level of assembly, the bulkier the complex will become, which will take longer time to pass through the matrix, and hence the bands show retarded patterns.

The truncated variants of the DNA hydrogels (DX0, PX0, TT0, TQ0, TP0, and TH0) show a minimal shift in their band patterns to the right side of the DNA ladders (Figure 2), due to their strand-directed hybridization under the influence of thermal annealing effects. All the network type DNA structures show significantly larger band retardations compared to their ssDNA counterparts, indicating the formation of intricate networks of DNA strands, which hinder their electrophoretic movements through the tight mess of the acrylamide gel.

**Figure 1:**
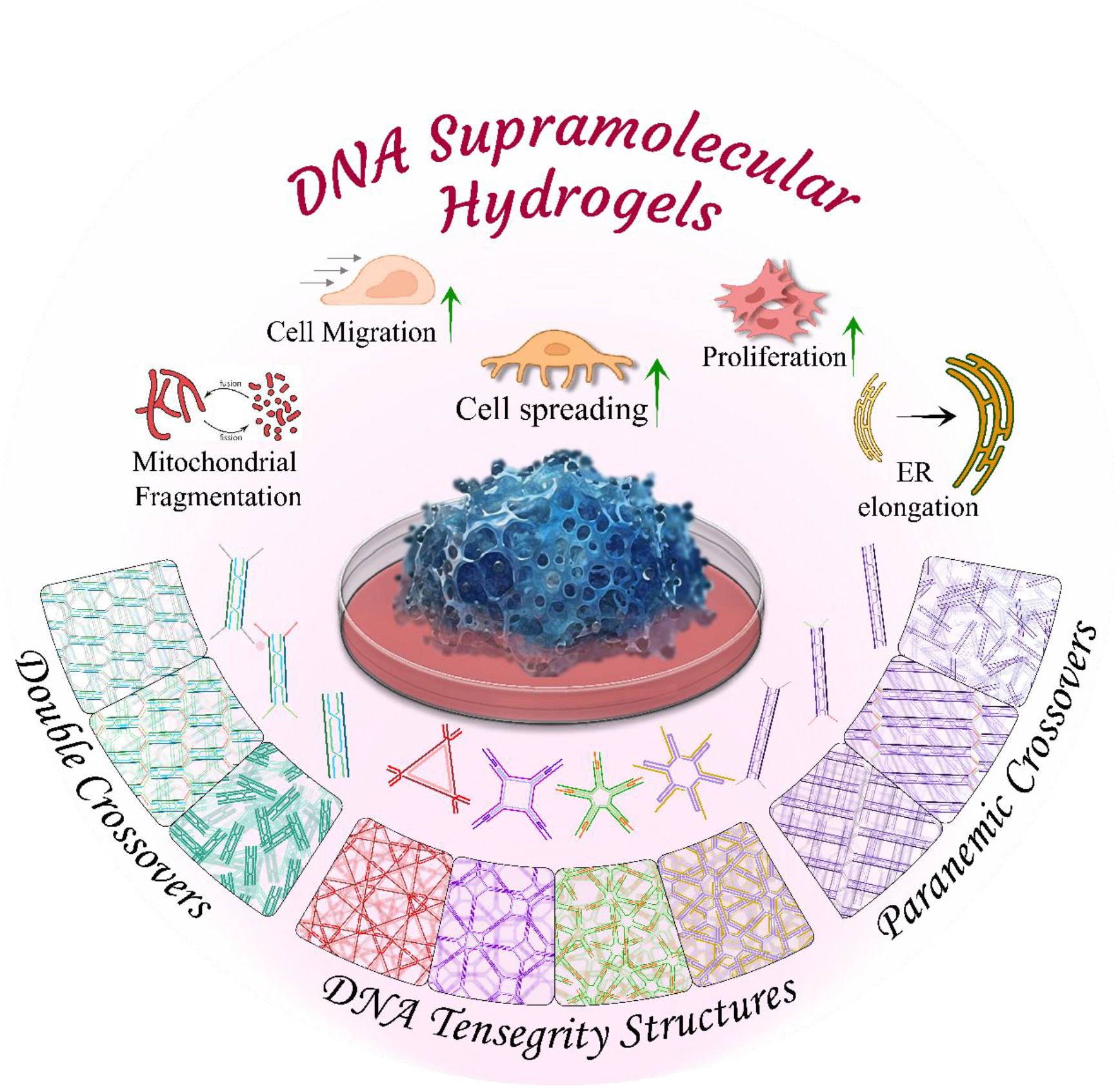
Graphical Illustration of customizable rigid DNA hydrogels with bespoke mechanical properties for modulating cellular behaviour of RPE1 cells in a mechanoregulated manner. Representation of free and polymerized DNA supramolecular assemblies of Double Crossovers (DX), Paranemic Crossovers (PX), and Tensegrity Structures.

**Figure 2:**
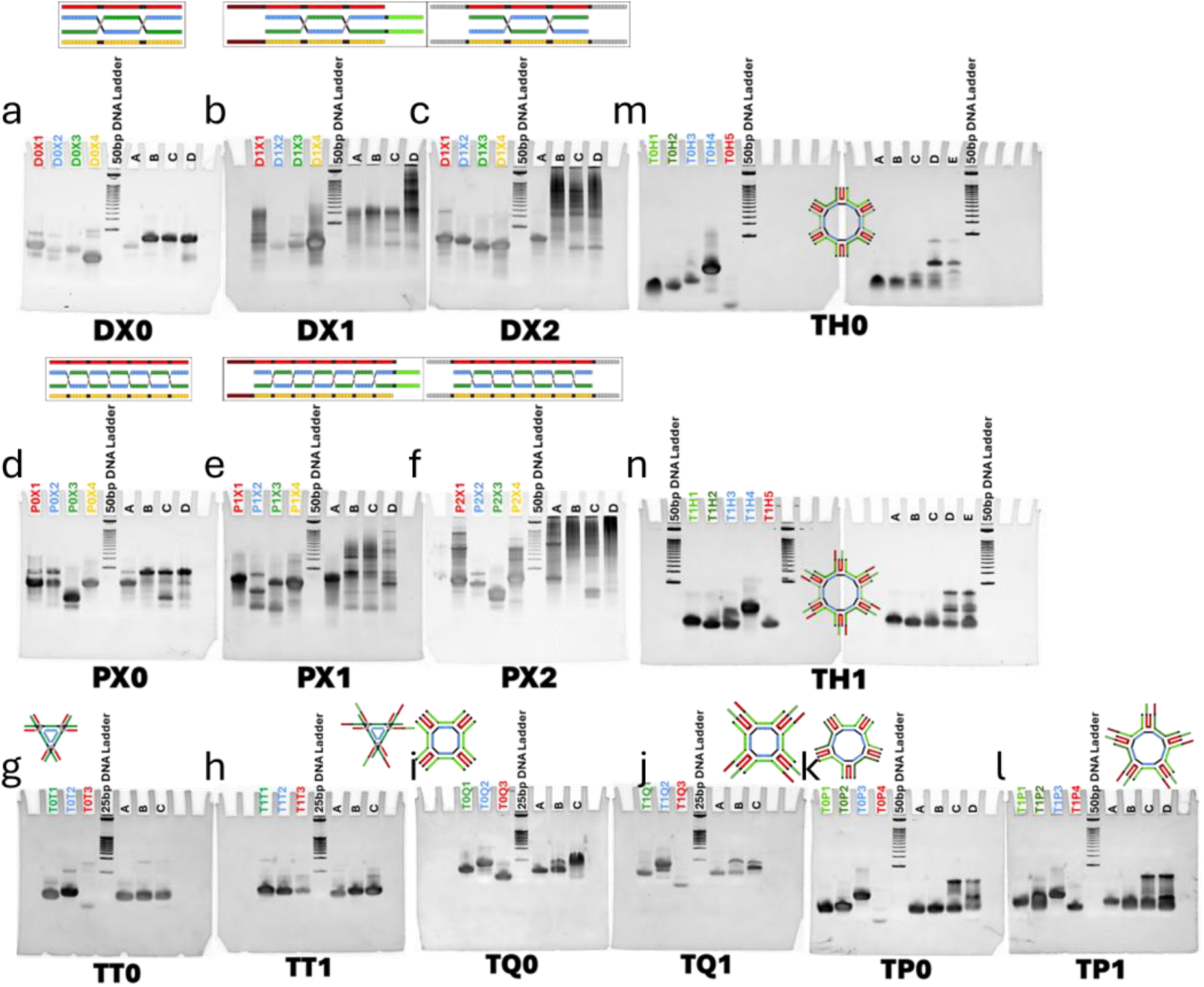
Figure shows EMSA band pattern of various rigid Hydrogels. On the left side of the DNA ladder, color-coded ssDNA is loaded, which forms higher order complexes in increasing order of assemblies on the right indicated by A,B, C, D, and E. Lane A, B, C, D and E denotes assembly of 1^st^ strand, 1^st^ and 2^nd^ strand, 1^st^, 2^nd^ and 3^rd^ strand, 1^st^, 2^nd^, 3^rd^, and 4^th^ strand and 1^st^, 2^nd^, 3^rd^, 4^th^ and 5^th^ strands, respectively.

Some DNA gels are notoriously escaping the combed wells of the 10% Native PAGE gels. This is observed especially in the case of the higher-order and highly branched DNA supramolecular structures. In these cases, only a minuscule amount of nucleic acids are able to traverse under the effects of the external electric field, which is demarcated by a faint or smear band. Through the visual inspection of band patterns alone, it is sufficient to say that the DX and PX hydrogel of type 2 network form complex and very thick hydrogels even at a loading volume of 3µL of sample.

#### Confocal-based imaging of DNA hydrogels

The lack of prominent and crisp band patterns in the EMSA assay, made it less reliable for assessing the supramolecular assembly and confirmation of DNA hydrogen network being formed. This ultimately comes down to the mechanical stiffness of the hydrogel. A probable solution would be to use a lower concentration of the cross-linked gel-based system for electrophoresis.

Another solution to assess the formation of nucleic acid-based DNA hydrogels is by fluorescently labelling them with DNA binding dyes to monitor the network morphology under liquid conditions. In this work, we have stained the DNA hydrogels with GreenR-a DNA intercalating dye with its excitation at 495nm and emission at 523nm. The GreenR bound DNA hydrogels, show a fluorescently labelled hydrogel morphology (**Figure 3**). The truncated DNA variants show no signs of Hydrogel network being formed. However, they aggregate together to form µm size, uniformly distributed DNA condensates. The probable reasons for the formation of condensates could be due to charge interactions between Mg^2+^ ions, which might partially neutralize the charges of the nucleotides, inducing liquid-liquid phase separation like behaviours. We also observe the size, and the intensity of the condensates increases with increasing arms of DNA tensegrity structures, indicating a significant involvement of branching architectures in determining fates of DNA condensates.

**Figure 3:**
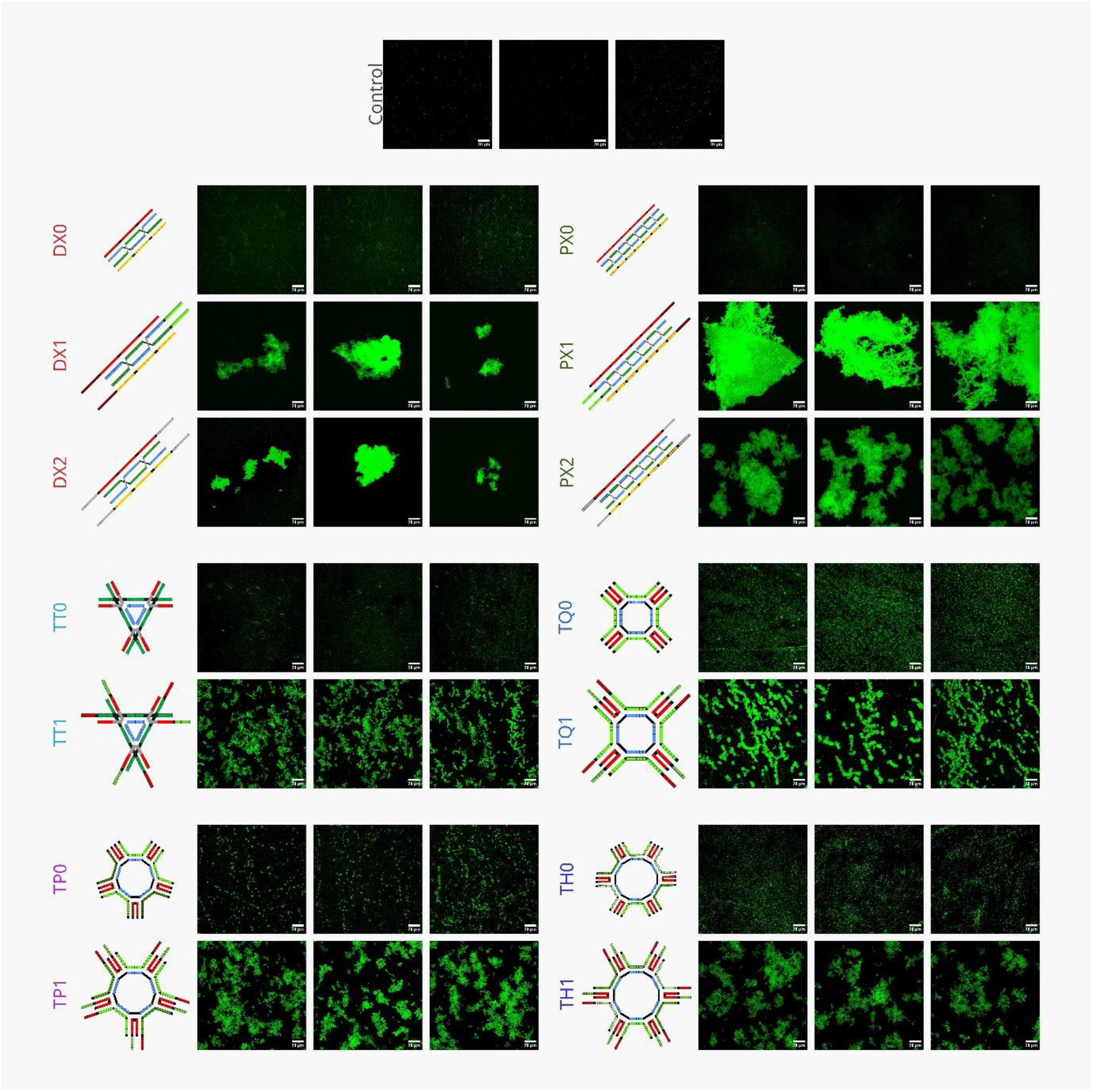
GreenR dye labelled DNA hydrogels imaged under confocal microscope reveals the network behaviour of the hydrogels. The truncated versions of DNA (DX0, PX0, TT0, TQ0, TP0, TH0) show no signs of branching morphology, indicative of hydrogel formations. The truncated DNA variants show µm scale aggreagates-like features indicative of formation of DNA condensates. The non-zero variants of DNA show diverse network forming hydrogels. The concentration of the DNA hydrogels was 1µM and stained with 0.01% GreenR dye. The scale bar is 20 µm.

**Figure 4:**
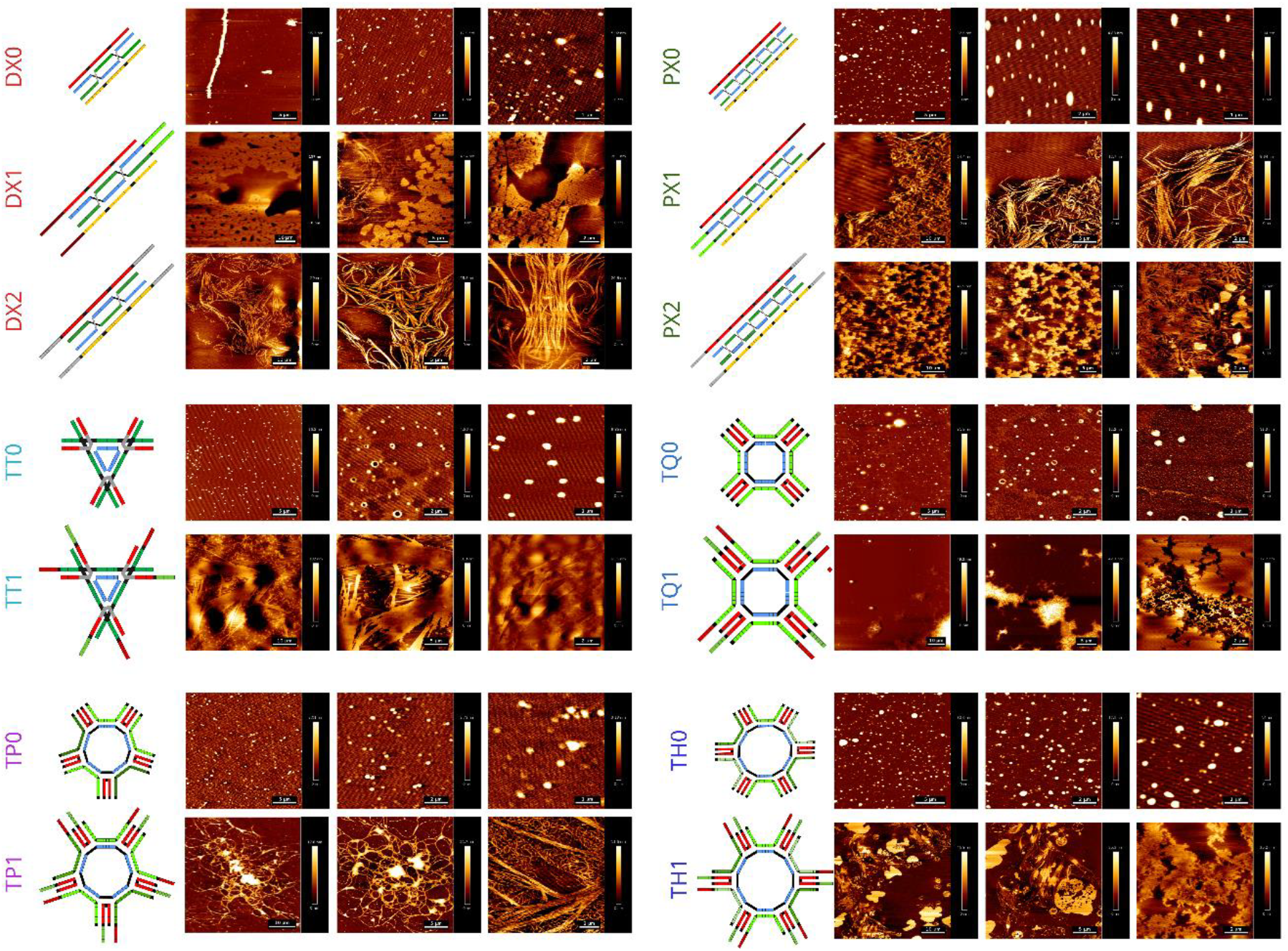
AFM-based morphological analysis of rigid DNA hydrogels. The scale bars of the left, middle and right images are 10µm, 5µm and 2µm, respectively.

The DX1 and PX1 motifs with specific sticky ends tailored to provide a directional assembly of motifs, offers a spatial directionality, and improved stiffness over palindromic sticky ends-type DX2 and PX2 counterparts. The nature of these sticky ends controls the hydrogel behaviour and network distributions at the nanometre scale, thereby offering precise control over engineering hydrogels catering to specific cell lines for advanced 3D tissue culture.

#### AFM-based morphology and rheology

When these DNA hydrogels are dried, they show an entirely different morphology. As expected, the truncated DNA motifs do not show any signs of assembly of interconnected chains. But incorporation of Sticky ends, enable then instantly to form hydrogels. The DX1 and PX1 variants having a directionally tuned sticky end, shows a mix of condensed granular and fibre like morphology. The polar nature of these sticky ends provides a spatial constraint over association of vacant DNA motifs, offering greater versatility, in terms of forms and functions. Random orientation of motifs, directed by either the sticky ends or lengthy core of the DNA motifs, give rise to fibrous hydrogel features, where each fibre consists of multiple strands of DNA assembled. Improving the stiffness of these motifs by incorporating tensile forces within these DNA strands, allows enhancement of mechanical properties, which is also reflected by tight packing of DNA fibres into thicker, longer and straight bundles of fibres (**TT1**). However, this tight packing is only visible when the core of the motif is small and compact. Increasing the size of the core by incorporating more strands of DNA destroys the compactness and induces flexibility to the fibres. The flexibility of the DNA fibres being packed is evident as we increase the number of arms from three arms in TT1 to six arms in TH1.

AFM-based Force Spectroscopic analysis revealed DNA hydrogel stiffness to be in the range of 50 kPa to 300 kPa. The truncated controls form condensates with higher Young’s modulus values compared to the blank control (containing NFW and MgCl_2_, at 35.7 kPa). Interestingly, the palindromic linker-based DX2 showed lower stiffness compared to DX1, however, the differences were not significant. But compared to PX0 (G’ = 50 kPa), the stiffness of PX2 (G 148.46 kPa) was threefold higher. The trend of rigidity showed an increase in the order of DX<PX< Tensegrity structures, supporting our main hypothesis. The stiffness, however, showed a decline in Pentameric tensegrity structures. This could be attributed to the stability of the pentameric junction in 3D space due to the non-symmetricity of junction-ends, affecting the overall stability of PX1 hydrogels, therefore, decreased stiffness.

Through the above results, the morphology and the rigidity of the hydrogel is dependent on several factors including the type of DNA motifs, number of branched structures available for network formation, length of the sticky ends facilitating the linkage, type of the sticky-end/ linkers, and the spatiodynamic core (motif junction) of the DNA hydrogels.

#### Cytotoxicity Assay of DNA Hydrogels

The cytotoxicity of synthesized DNA hydrogels was thoroughly tested at three different concentrations of 25 µM, 50 µM, and 100 µM, respectively. At lower concentrations of 20 µM DNA Hydrogels, the RPE1 cells showed slight favourability by showing an increase in the cell populations, as indicated by an increase in the absorbance of the formazan crystals thus formed during MTT assay, compared to our control, i.e., poly-L-lysine treated RPE1 cells. At higher concentrations of 50 µM (**Figure 5**), the cellular viability of RPE1 cells with respect to the treatment of the rigid DNA hydrogels was more pronounced, compared to the PLL treated group. Interestingly, in the Double crossover hydrogel group, the DX0 group, which does not form hydrogel network, shows comparable cell growth, as compared to the highest growth observed in DX1. Comparing all the candidates of DX type, the change in cell viability of RPE cells is non-significant, nonetheless these show threefold cellular viability compared to PLL. As earlier demonstrated, the truncated control groups of DNA hydrogels actively form condensate-like morphology, and are uniformly distributed across the surface, we propose that these truncated DNA controls provide miniaturized hotspots of scaffold-like environment, thereby maximizing the cellular viability of RPE1 cells compared to their network-forming counterparts.

**Figure 5:**
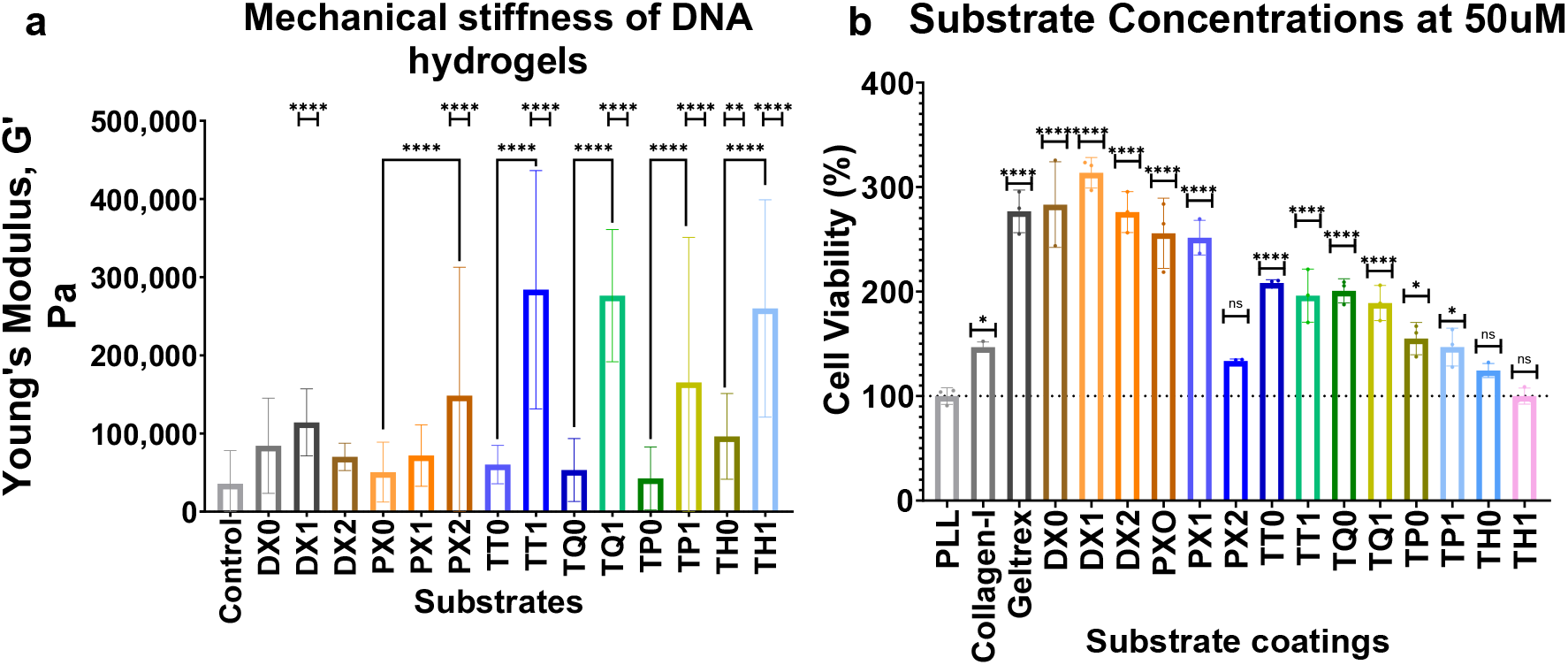
(a) Graph showing mechanical stiffness of DNA hydrogels through Force Spectroscopy using Bio-AFM instrument. (b) Cell viability of RPE 1 cells grown over DNA hydrogels at 50uM concentrations. Simple One-way ANOVA was used to compare the results. **** signifies p<0.0001.

At the highest concentration of 100 µM, DNA hydrogels show even higher cellular viability response compared to lower DNA HG concentration levels. On average, all DNA hydrogels, whether truncated or networking-type, show four-fold cellular viability compared to PLL alone. The double-crossover type DNA systems provide the maximum cellular viability response. From the aspects of complexity and crossovers, it is expected that tensegrity structures have higher mechanical stiffness compared to DX and PX type hydrogels. And from the behaviour of RPE 1 cells, it is certain that RPE 1 cells appreciate hydrogels of a comparatively softer stiffness profile.

### c. Cellular adhesion and spreading

The RPE 1 cells are epithelial cells with generally spindle-shaped morphology on flat substrates, like glass or plastic culture dishes. Stiffer substrates promote cytoskeletal organization and distribution, which applies traction forces over the cellular focal adhesion sites and thereby triggers the cells to contract. This process of stiffness-induced contraction allows the cells to become elongated and spindle-like. Another effect of ECM stiffness allows the cells to become enlarged, compared to constricted growth over flat or soft substrates.

Amongst the controls that we have used in this study, PLL consists of a layered aggregation of poly-L-lysine, which coats the substrate with a soft and fibrous coating. Collagen I is a highly purified component of ECM and defines the stiffness of the tissues, including bones and cartilage. Collagen I cannot entirely represent the ECM alone. Geltrex is a commercially available cocktail of ECM components, including laminin, Collagen IV, entactin, heparin sulphate proteoglycans, etc. The Geltrex can therefore supplement the RPE1 cells with overall growth conditions as can be expected in an *in-vivo* site. The last control is a glass coverslip, which provides a micron-level surface for cells to anchor and grow in the presence of cell-culture grade media.

**Figure 6** shows single-cell images of RPE 1 cells cultured over various substrates of varied stiffness. Upon visual inspection, it is evident that the actin organization and assembly are slightly reduced in softer substrates like PLL, collagen I, and Geltrex compared to DNA hydrogel samples. Actin staining gives a rough estimation of cell shape, size, and area. Quantification of these samples reveals that DNA hydrogels enhance the cells’ spreading potential by modulating actin reorganization. We observed an increase in cell area for the cells grown over truncated control groups compared to the intact hydrogel network groups at lower mechanical stiffness levels. This phenomenon is more pronounced at stiffer hydrogels like TT1, TQ1, TP1, and TH1. We also observed that the change in cellular area for truncated and non-truncated DNA motifs is not significant. This observation could be explained by the specific mechanoelastic preference of RPE 1 cells. Nanoindentation studies by Amela et. al., for measuring the stiffness of cortical and nuclear cells of the eyes, reveal that the stiffness of these cells was within the range of 0.1 kPa to ∼20 kPa (Hozic *et al*., 2012). Another study highlighted the stiffness of corneal cells to be in the range of 7.5 kPa to 109.8 kPa (Last *et al*., 2012). These evidences have a direct correlation with the stiffness of the DNA hydrogels. The increase in cellular area keeps increasing along with an increase in the stiffness of DNA hydrogels till PX2 hydrogels’ Young modulus values, which is close to the innate stiffness of RPE1 cells. Moving higher in terms of scaffold stiffness resulted in a decreasing trend in cellular area. Furthermore, the stiffer hydrogels like TT1, TQ1, TP1, and TH1 offer an unfavourable microenvironment to the RPE1 cells, which is indicated as a decline in cell area. However, the condensates of stiffer DNA motifs provide the necessary mechanical forces to the RPE1 for their optimal growth and development, which is reflected in terms of higher cellular area compared to hydrogels of DX and PX junctions (**Figure 6.b**).

The nucleus of RPE 1 also shows a hydrogel stiffness-dependent mechanomodulatory response. RPE1 cells grown over hydrogels are significantly larger (∼ 2x larger than control) than the PLL group (**Figure 6.b**).

**Figure 6:**
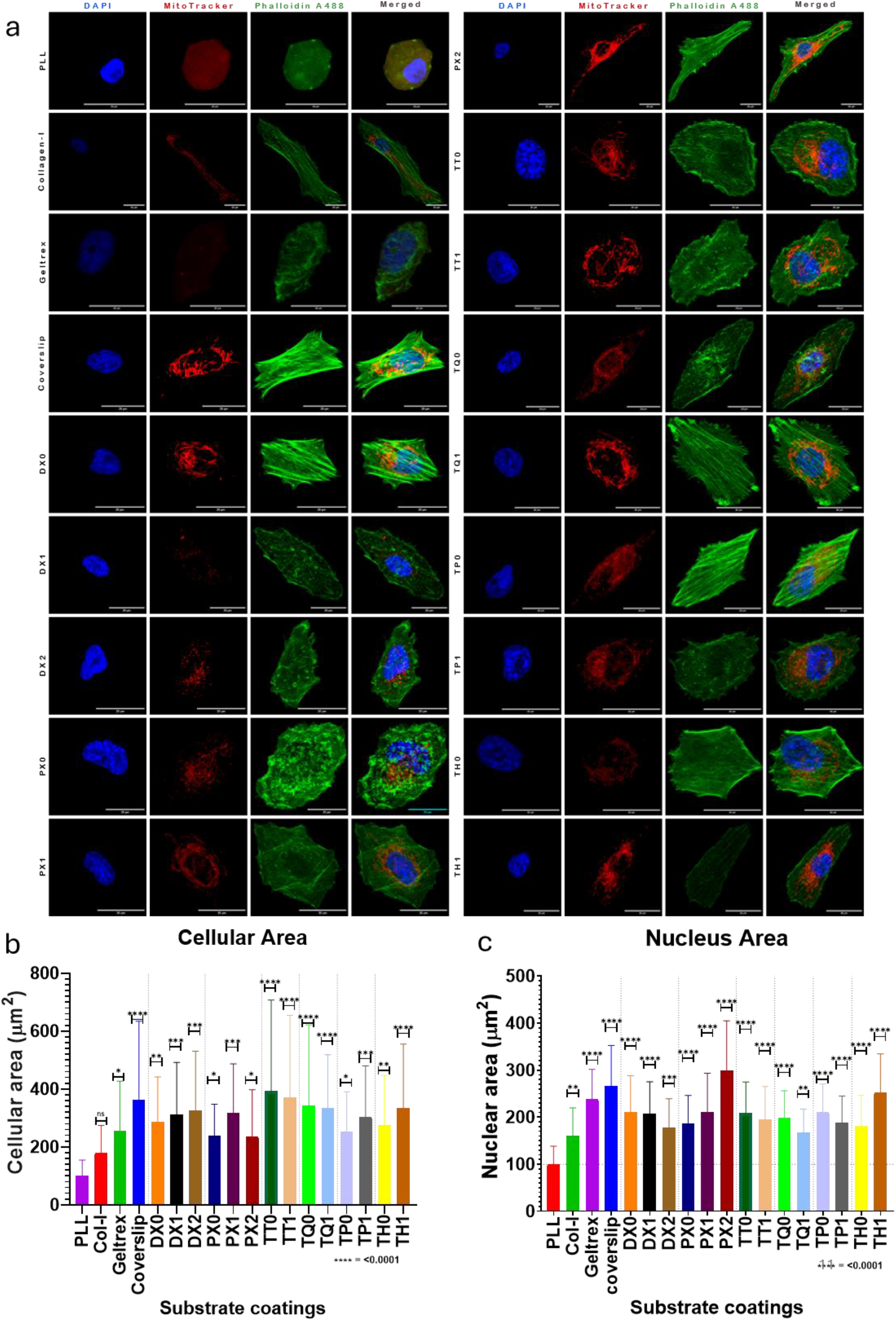
Cellular area and Nucleus area perturbations in RPE1 cells’ morphology due to substrate stiffness. (a) Single-cell images of actin, nucleus, and mitochondrial expression and distribution. (b,c) Quantified and analyzed fluorescence data of actin and nucleus show stiffness-dependent cell and nucleus area enlargement in RPE1 cells. The scale bar is 20 µm. A simple one-way ANOVA was performed to compare the graphs. **** indicates p-value <0.0001

### d. Expression pattern and distribution of Mitochondria and Endoplasmic reticulum

ECM stiffness regulates the cytoskeletal organization, which affects the cell morphology and organelles. Increased cellular area and proliferation upon growth on DNA hydrogels indicate a metabolically active state of the cell, which creates a pressure on normal homeostatic states of the cellular organelles. Faster rate of replication coupled with enlarged cellular morphology is often associated with rapidly regulating the cell’s organellar distribution and organization. Endoplasmic reticulum and Mitochondria are two of the most widely studied cellular organelles, whose distribution and regulations have a strong correlation with ECM stiffness.

Activation of mechanotransducers like Piezo1 triggers Ca2+ influx via activation of ERK signalling pathways (O’Rourke *et al*., 2002; Ke *et al*., 2023). This led to mitochondrial fission, with reduced tubular networks and reduced fluorescence signal of the MitoTracker DeepRed dye. As a result, we see a diminution of fluorescence signal in correspondence with an increase in scaffold stiffness levels. Increased cellular proliferation diverts the cells’ attention from oxidative phosphorylation to glycolysis for instantaneous energy supply to meet the needs of actively dividing cells (Singh *et al*., 2024; Wang *et al*., 2025). Fragmented mitochondria are easily accessible for faster ATP turnovers during cellular division, which adds up in the support of our findings.

The changes in ER expression levels seem likely to stem from mechanotransduction-driven ER structural remodelling and stress response attenuation (Gruber *et al*., 2023; El Yousfi *et al*., 2025). Stiffer hydrogels produce cellular tractions through acto-myosin contractions, resulting in elongation of the cell, which otherwise remain circular (**Supplementary Figure 14**). These tensions are transmitted to the ER networks through mechanotransducers like nesprins, causing stretching and expansion of ER tubules. This change is perceived as an increment in ER tracking fluorophores’ signals.

We observed a positive correlation between increased MitoTracker expression patterns and hydrogel stiffness values in the range of 100 kPa. Once the scaffolds cross this 100 kPa threshold, the MitoTracker expression pattern decreases. Closer inspection of the single-cell images reveals that the fragmentation profile of mitochondria increases with an increase in stiffness of DNA hydrogels (**Figure 7.b**). This could explain the increased fluorescence of fragmented mitochondria distributed throughout the cells. At higher stiffness values, >100 kPa, the RPE 1 cells show a decrease in cellular area and proliferation, correlating positively with a decrease in mitochondrial absorbance and increased filamentation of the mitochondrial networks (**Figure 7**).

**Figure 7:**
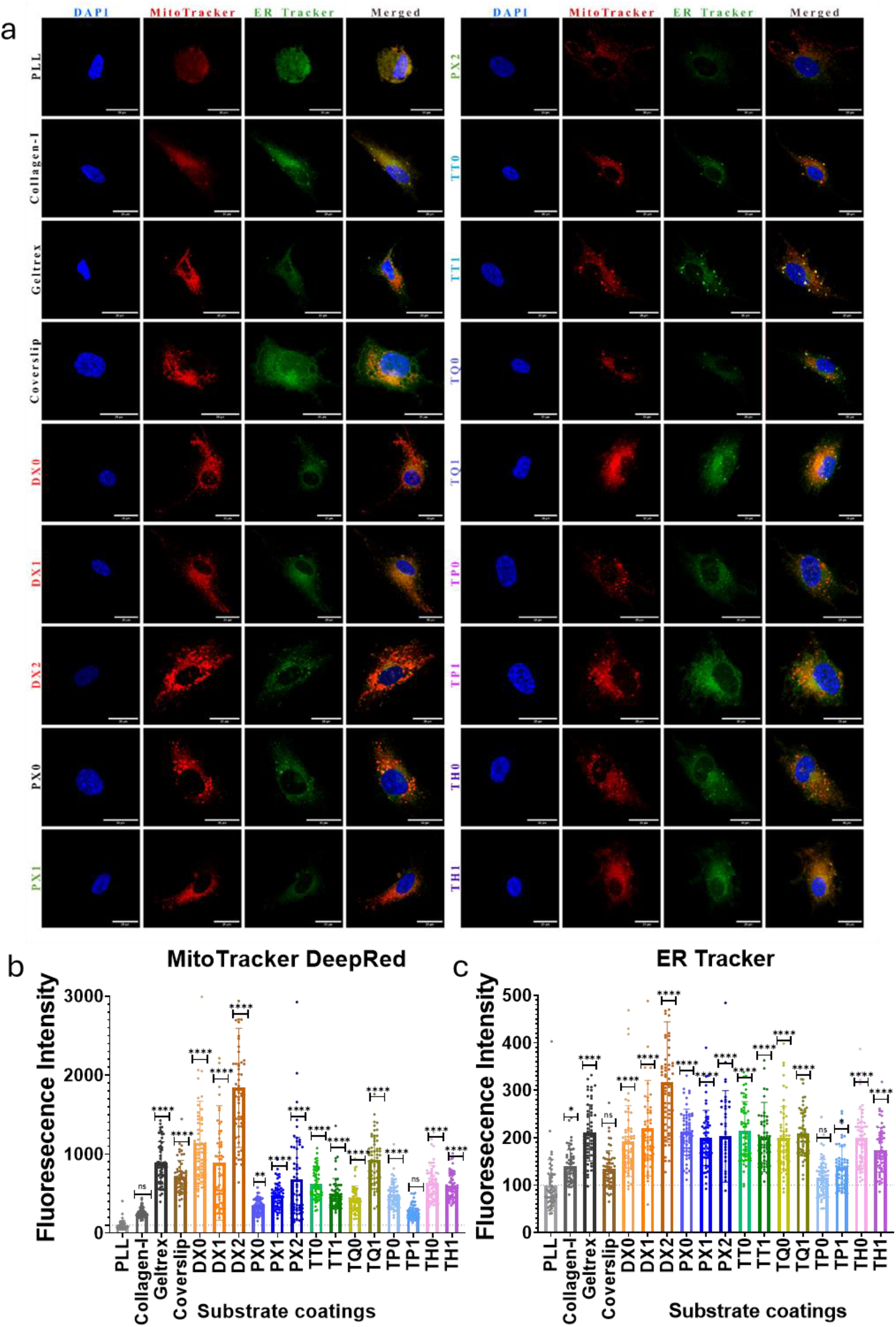
Dynamics of Mitochondrial and Endoplasmic Reticulum are mechanoregulated via rigid DNA hydrogels. (a) Single-cell images of RPE1 cells cultured over DNA hydrogels and controls. The cells are stained with DAPI (nucleus, blue), MitoTracker DeepRed (Mitochondria, red), and ER Tracker-BODIPY-FL (ER, green). Qunatification of Fluorescence images of (b) MitoTracker DeepRed and (c) ER tracker-BODIPY FL stained RPE 1 cells. The scale bars are 20µm. **** signifies p-value <0.0001.

Inversely, the ER tubules get elongated with increased cellular traction as a result of mechanotransductionary regulation from DNA hydrogels. Below the threshold of 100 kPa Young’s moduli, the elongation in ER directly follows the trends of increases mechanoelasticity of the substrate, however after crossing the threshold the expression of ER tracker drops significantly (**Figure 7.c**).

## III. Materials & Methods

The ssDNA strands (**Supplementary Table 1**) used in this study were purchased at 0.2µM production scale from Merck (Sigma-Aldrich), followed by desalting purification. The lyophilized ssDNA strands were dissolved in Nuclease-Free Water (NFW) at 72°C for 60 minutes and stored at 4°C until further use. Moviol, Triton-X, DNA loading dye, Phalloidin-Alexa 488, and DAPI were purchased from Merck (Sigma-Aldrich). We bought APS (ammonium persulfate), EtBr (ethidium bromide), Triton-X, TEMED (tetramethylethylenediamine), paraformaldehyde, cell culture dishes, and flasks for adherent cells (treated surface) from Himedia. We procured DMEM (Dulbecco’s Modified Eagle’s Medium), FBS (Fetal Bovine Serum), Pen-strep (penicillin™streptomycin), and 0.25% trypsin™ethylenediaminetetraacetic acid (Trypsin-EDTA) from Gibco. GeNei provided TAE (Tris-acetate-EDTA) and 30% Acrylamide / Bisacrylamide solution (29:1). The MgCl_2_ (Magnesium chloride), NaCl (Sodium Chloride), KCl (Potassium Chloride), Na_2_HPO_4_ (disodium hydrogen phosphate), and KH_2_PO_4_ (potassium dihydrogen phosphate) were purchased from SRL India and Santa Cruz Biotech. Gene to protein provided us with GreenR-DNA gel stain. Hi-grade Mica sheets (grade 2) were purchased from Ted Pella, and microslides were procured from Bluestar.

### 1. Design and synthesis of DNA hydrogels

Through our previous work, we demonstrated the design, synthesis, characterization, and applications of soft DNA hydrogels made up entirely of DNA duplex-inspired architectures. In this work, we try to explore other DNA supramolecular motifs to form rigid/ mechanically stronger complexes to form hydrogels. Here, we have demonstrated fourteen unique DNA supramolecular Complexes, where six complexes have truncated blunt ends acting as our intrinsic controls, making them incapable of forming hydrogel networks, while the remaining complexes have sticky ends, enabling them to form geometrically defined intricate networks.

The ssDNA strands are designed through a rational approach while relying on the Watson-Crick base-pairing rule. The strands were enriched with high GC content, where the binding was expected/ programmed to be higher (at the ends) as compared to the availability of AT-rich regions (cross-over sites), where flexibility of the strands was to be expected. Depending on the complexity of the supramolecular design, the number of ssDNA strands varies. For example, to form DX and PX structures, four strands of ssDNA were utilized in equimolar (1:1:1:1) concentrations to form the complex. The complexes like TTs, TQs, TPs, and THs have the stoichiometric ratios of ssDNA in 3:1:3 (Three ssDNA in TT); 4:1:4 (Three ssDNA in TQ); 3:2:1:5 (four ssDNA in TP); and 2:2:2:1:6 (five ssDNA in TH), respectively.

Based on the envisioned complex structure design, a 2D framework was first made over the scadNANO website (https://scadnano.org/). Based on the above 2D framework (DX, PX, TTs, TQs, TPs, and THs), the sequences were rationally integrated in an iterative manner to maximize thermodynamic stability while minimizing unwanted crossovers and potential to form primer-dimers, secondary loop structures. The flowmap of the design process is demonstrated via the flowchart attached in the supplementary (**Supplementary Figure 1**). The ssDNA sequences were then fed to the open-source RNAStructures web server (https://rna.urmc.rochester.edu/RNAstructureWeb/) to predict secondary structure in a pairwise-assembly mode using the DuplexFold Algorithm. The Pair-wise Gibbs free energy values were used to scrutinize the sequences iteratively to arrive at the final refined sequences.

These refined sequences were then synthesized and purified. These sequences, as per their specific architecture type, are pooled in a PCR reaction tube, in the presence of MgCl_2_ ions, followed by a thermal annealing protocol as reported earlier (Singh and Bhatia, 2022; Singh *et al*., 2024; Singh, Singh and Bhatia, 2024). After gradually cooling down the hydrogels, they are stored at 4°C until further use.

### 2. Characterization of DNA hydrogels

The hydrogels formed by various ssDNA assemblies are similar in nature yet different in terms of size, shape, structure, and morphology, which makes them excellent for programming cellular behaviours. These DNA hydrogels are characterized by the following techniques.

#### i. Electrophoretic Mobility Shift Assay (EMSA)

The EMSA is usually performed over a dense gel system, usually with the help of electrophoresis. In our case, we used a 10% Native PAGE gel prepared using a 30% acrylamide/bisacrylamide solution (29:1), crosslinked with 10% APS and 10µL TEMED in 5x TAE solution. The freshly prepared DNA Hydrogel samples (2µL) were then mixed with 1µL of 6x DNA-loading dye, followed by loading them into the wells for electrophoretic separation. The EMSA was carried out in ice-cold TAE buffer to minimize the temperature generated during electrophoresis. The powerpack was programmed to run for 90 minutes at 90V for efficient separation. The bands of the separated DNA hydrogel samples were stained with EtBr for 15 minutes under gentle shaking, followed by imaging through the Bio-Rad gel-documentation systems.

#### ii. Confocal-based imaging of DNA hydrogels

10 µL of freshly prepared DNA hydrogels (10 µM) were mixed with 1uL of 1/1000 dilution of GreenR intercalating dye (Ex/Em =495nm/ 523nm). The mixed samples were then directly loaded on a freshly cleaned glass slide, followed by covering the sample with a thin coverslip. The prepared slides were then taken for confocal microscopy-based imaging of GreenR-labelled DNA hydrogels.

#### iii. AFM-based morphology of DNA hydrogels

The freshly prepared DNA samples were directly drop-cast over a cleaved mica sheet. The samples were stored in a vacuum desiccator for air drying. Once the samples were completely dried, the hydrogels’ morphological features were imaged using Bruker’s NanoWizard Sense+ BioAFM instrument. The imaging was carried out in tapping mode while using a Scanasyst-Air tip with a 2nm tip diameter, operating at 75 kHz oscillating frequency in Direct drive mode. The samples were imaged in a 512*512-pixel format.

AFM-based Force spectroscopy (AC-mode) was performed using ScanAsyst-Air tips (Φ = 2nm, k = 0.35 N/m) over Bruker’s NanoWizard sense plus instrument. The scan was performed in a relatively closed-loop mode, measuring a total of 256 points per area (16 points * 16 points) while keeping the force of the interacting tip constant. The force maps were plotted using vertical deflection (nN, Y-axis) and measured Heights (µm, X-axis) and batch processed by performing an elastic fit using the Hertz/Sneddon model. The output generates a histogram of Young’s modulus of 16 bins of the sample. The average of six such histograms was compiled to represent the final graph (**Figure 5.a**)

#### iv. *In-vitro* Assays

##### Cytotoxicity Assay

The effect of DNA hydrogels on RPE 1 cells was analyzed by performing an MTT-based cytotoxicity assay. In a 96-well plate, all fourteen DNA hydrogel samples with varying concentrations, along with controls, were pre-coated and kept for overnight incubations. After which, cells were seeded at a seeding density of 10,000 cells/ well. The cells were grown over the substrates for 24 hours. The old media was decanted and washed with 1x PBS before adding MTT reagent at a concentration of 0.5mg/mL. The MTT added culture plates were then kept for incubation at 37°C under dark conditions. After four hours of incubation, the MTT reagent was removed, and DMSO was added, followed by gentle rocking to dissolve the formazan crystals. The

##### Cell adhesion and spreading

The RPE1 cells were cultured till 90% confluency, followed by seeding over 10mm coverslips coated with different substrates at a seeding density of 10,000 cells/ well. The substrate samples included PLL, Collagen-I, Geltrex, and DNA hydrogels at 50 µM concentrations. After 24 hours of incubation, the cells were washed off with PBS, followed by fixing under 4% PFA conditions for 15 minutes. The remaining PFA was washed off with PBS again after the incubation, followed by sequential staining of RPE 1 cells with 1µg/mL MitoTracker deep red in PBS for 15 minutes, following the Phalloidin-Alexa488 (in 0.1% Triton X-100) staining protocol as reported in our previous work (Singh *et al*., 2024).

##### Mitochondrial and ER staining

For the study of organellar distribution, the staining was performed after 4% PFA fixation, followed by PBS washing. The staining solution was prepared by adding 1µg/mL of Mitotracker DeepRed, and 1µL of 1 mM ER tracker Green-BODIPY FL in 1 mL PBS. In place of PBS, serum-free media could also be used. All the staining processes post-fixation were performed by placing the cells over an ice block (4°C refrigerator) to minimize the unbinding of the stains from the target site, protein denaturation, and signal losses.

## Conclusions

Our research demonstrates that DNA supramolecular hydrogels offer unprecedented control over mechanical properties, with higher branching complexity correlating to increased mechanical stiffness. The incorporation of DX, PX, and tensegrity structures along with specific linker types allowed us to make a range of mechanically tuneable DNA hydrogels of specific stiffness and network dynamics. We demonstrated a whopping increase in polymeric stiffness by incorporating tensegrity structures, where the mechanical stiffness reached 283.95 kPa of polymeric stiffness.

These hydrogels also regulate the cellular proliferation in a concentration-dependent manner, where higher concentrations boost the proliferation of the cells up to an eightfold change compared to the PLL controls. These hydrogels regulated the cellular behaviour, including their cellular adhesion, spreading, proliferation, shape, and organellar dynamics, including mitochondrial fragmentation and ER stress attenuation. We demonstrated the mechanomodulatory effects of mechanically tuneable hydrogels for RPE1 cell culture and development.

RPE1 cells show an increment in cellular area in hydrogels with < 100 kPa Young’s moduli. This observation is coordinated with an active remodelling of ER reticulum and Mitochondrial states in a stiffness-dependent manner. RPE1 cells start showing signs of reduced cell area coupled with mitochondrial filamentation and reduced ER signals upon exposure to hydrogels with G’> 100 kPa.

By systematically investigating the relationship between DNA structural complexity and resulting mechanical properties, we have established a framework for the rational design of DNA hydrogels with precisely tuned stiffness and geometric features. The ability to program both the structure and function of these DNA hydrogels through sequence design paves the way for significant advances over conventional hydrogel systems, offering new possibilities for creating instructive cellular microenvironments for precise modulation of cell behaviour and tissue development.

### Future directions

The ability to control both the mechanical properties and cellular responses, through DNA hydrogel design, opens numerous avenues for application in tissue engineering and regenerative medicine. By matching the desired mechanical properties of the target cells/ tissues by promoting specific cellular responses through mechanoregulation, these hydrogels could be scaled up as instructive scaffolds that not only support cell growth, but also actively guide the tissue development and functions. The ease of customizability and functionalization of DNA scaffolds offers a lucrative opportunity to further modulate specific cellular responses.

Addition of stimulus-responsive motifs like G-quadruplexes, i-motifs, CpG regions, hairpin loops, and aptamers extends the reach of this study even beyond. Incorporation of these unique motifs can be used to make the DNA hydrogels dynamically responsive to real-time changes in temperature, pH, solute/solvent concentrations, etc. This approach would more closely mimic the dynamic nature of the natural ECM, which undergoes continuous remodelling during tissue development and regeneration.

Also, the precision of DNA network assembly offers the possibility of spatio-temporally ligating molecules of differentiation, leading to a robust platform for studying head-tail polarity or more complex phenomena of developmental biology. The programmability of DNA hydrogels also opens avenues for personalized medicines in the future, where the mechanical properties of these hydrogels could be tuned according to specific patients’ needs. By integrating patient-derived cells with custom-designed DNA hydrogels, it would become possible to create more efficient disease models for testing drugs and personalized therapeutic regimens.

We have demonstrated pure DNA-junction-based hydrogels with customizable mechanoelastic properties. In the future, we would like to explore photodynamic regulation of DNA hydrogels conjugated with photosensitizers for developing photodynamic drug-delivery scaffolding systems for cancerous or tumor models.

## Author Information

Ankur Singh-Department of Biological Sciences and Engineering, Indian Institute of Technology Gandhinagar, Gandhinagar, Gujarat 382355, India.

Dhiraj Bhatia – Department of Biological Sciences and Engineering, Indian Institute of Technology Gandhinagar, Gandhinagar, Gujarat 382355, India.

## Authors’ contribution

D.B. conceived the idea, and A.S. and D.B. planned the experiments. A.S. did the synthesis, characterization by EMSA, Bio-AFM, confocal microscopy, and *in-vitro* cell culture experiments. A.S. analysed and processed all the data and was cross-checked by all mentors. All the authors contributed to writing and reading the manuscript draft.

## Acknowledgements

We sincerely thank all the members of D.B.’s group for critically reading the manuscript and for their valuable feedback. A.S. thanks PMRF for funding his Ph.D. We acknowledge the central instrumentation facilities at IIT Gandhinagar. The work in host laboratories is funded by ANRF-CRG, GSBTM and MoES-STARS grants.

## Ethics declarations

### Competing interests

The authors declare no competing interests.

## Notes

### Competing Interest Statement

The authors have declared no competing interest.

