## Supporting information for "DNA cross-over motifs based, programmable supramolecular hydrogels for mechanoregulatory effects of cellular behaviour and cytoskeleton reorganization"

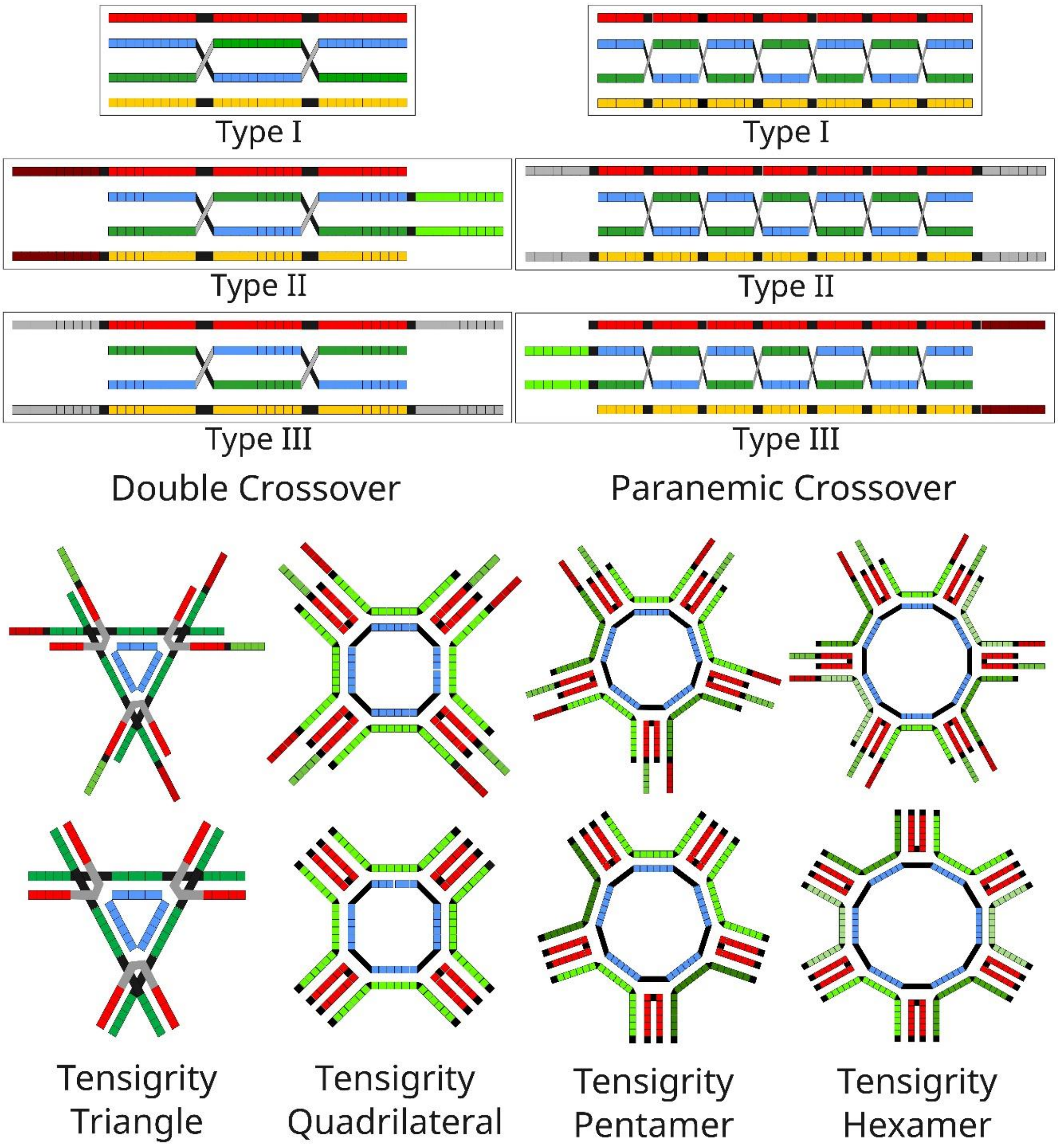

*Figure 1: Graphical illustration of DNA motifs with DX, PX, and Tensegrity structures. Each individual box indicates the position of a nucleotide, except black boxes, which indicate gaps. The linker types are color-coded for palindromic (grey color, non-specific), and non-palindromic (red/green, directional assembly)*

| Sr. No | HG Type | Name | Sequence | Sequence length | GC% | Ratio |
|---|---|---|---|---|---|---|
| 1 | DX0 | D0X1 | GGTTCAGATTAAGTCTCTAATTTAGTTCGC | 30 | 36.6 | 1 |
| 2 | | D0X2 | GCGAACTAAAGCTATACACGAATCTGAACC | 30 | 43.3 | 1 |
| 3 | | D0X3 | GATCAACCGACGATATGTGCCCACCAAACC | 30 | 53.3 | 1 |
| 4 | | D0X4 | GGTTTGGTGGATCACATCGCTCGGTTGATC | 30 | 53.3 | 1 |
| 1 | DX1 | D1X1 | CGTGCAGCGC GGTTCAGATT AAGTCTCTAA TTTAGTTCGC | 40 | 47.5 | 1 |
| 2 | | D1X2 | GCGCTGCACG GCGAACTAAA GCTATACACG AATCTGAACC | 40 | 52.5 | 1 |
| 3 | | D1X3 | GCGCTGCACG GATCAACCGA CGATATGTGC CCACCAAACC | 40 | 60 | 1 |
| 4 | | D1X4 | CGTGCAGCGC GGTTTGGTGG ATCACATCGC TCGGTTGATC | 40 | 60 | 1 |
| 1 | DX2 | D2X1 | GCGCTAGCGC GGTTCAGATT AAGTCTCTAA TTTAGTTCGC CGCGATCGCG | 50 | 54 | 1 |
| 2 | | D2X2 | GCGAACTAAA GCTATACACG AATCTGAACC | 30 | 43.3 | 1 |
| 3 | | D2X3 | GATCAACCGA CGATATGTGC CCACCAAACC | 30 | 53.3 | 1 |
| 4 | | D2X4 | GCGCTAGCGC GGTTTGGTGG ATCACATCGC TCGGTTGATC CGCGATCGCG | 50 | 64 | 1 |
| 1 | PX0 | P0X1 | GTGGT ATCAT CAATG CTATG TGTAG GCTTA GACCT | 35 | 42.8 | 1 |
| 2 | | P0X2 | AGGTC CGAAA CTACA CAATT CATTG CCGAA ACCAC | 35 | 45.7 | 1 |
| 3 | | P0X3 | CCGTC TAAGC TGCCA CATAG ACAGC TGTAT CACAG | 35 | 51.4 | 1 |
| 4 | | P0X4 | CTGTG TTCGG GCTGT AATTG TGGCA TTTGC GACGG | 35 | 54.2 | 1 |
| 1 | PX1 | P1X1 | CGTGCAGCGC GTGGT ATCAT CAATG CTATG TGTAG GCTTA GACCT | 45 | 51.1 | 1 |
| 2 | | P1X2 | GCGCTGCACG AGGTC CGAAA CTACA CAATT CATTG CCGAA ACCAC | 45 | 53.3 | 1 |
| 3 | | P1X3 | GCGCTGCACG CCGTC TAAGC TGCCA CATAG ACAGC TGTAT CACAG | 45 | 57.7 | 1 |
| 4 | | P1X4 | CGTGCAGCGC CTGTG TTCGG GCTGT AATTG TGGCA TTTGC GACGG | 45 | 60 | 1 |
| 1 | PX2 | P2X1 | GCGCTAGCGC GTGGT ATCAT CAATG CTATG TGTAG GCTTA GACCT CGCGATCGCG | 55 | 56.3 | 1 |
| 2 | | P2X2 | AGGTC CGAAA CTACA CAATT CATTG CCGAA ACCAC | 35 | 45.7 | 1 |
| 3 | | P2X3 | CCGTC TAAGC TGCCA CATAG ACAGC TGTAT CACAG | 35 | 51.4 | 1 |
| 4 | | P2X4 | GCGCTAGCGC CTGTG TTCGG GCTGT AATTG TGGCA TTTGC GACGG CGCGATCGCG | 55 | 63.6 | 1 |
| 1 | TT0 | T0T1 | GGCCA GTACGG TGCCG | 16 | 75 | 3 |
| 2 | | T0T2 | TAC CCGTAC CCGTAC CCG | 18 | 66.6 | 1 |
| 3 | | T0T3 | CGGCA TGGCC | 10 | 80 | 3 |
| 1 | TT1 | T1T1 | AGCGCT GGCCA GTACGG TGCCG | 22 | 72.7 | 3 |
| 2 | | T1T2 | TAC CCGTAC CCGTAC CCG | 18 | 66.6 | 1 |
| 3 | | T1T3 | AGCGCT CGGCA TGGCC | 16 | 75 | 3 |
| 1 | TQ0 | T0Q1 | GTCAGA ATCGGA TGCCAG | 18 | 55.5 | 4 |
| 2 | | T0Q2 | GAT TCCGAT TCCGAT TCCGAT TCC | 24 | 50 | 1 |
| 3 | | T0Q3 | CTGGCA TCTGAC | 12 | 58.3 | 4 |
| 1 | TQ1 | T1Q1 | GCTCA GTCAGA ATCGGA TGCCAG | 23 | 56.5 | 4 |
| 2 | | T1Q2 | GAT TCCGAT TCCGAT TCCGAT TCC | 24 | 50 | 1 |
| 3 | | T1Q3 | TGAGC CTGGCA TCTGAC | 17 | 58.8 | 4 |
| 1 | TP0 | T0P1 | GTCAGA GCCATA TGCCAG | 18 | 55.5 | 3 |
| 2 | | T0P2 | GTCAGA CTAGCC TGCCAG | 18 | 61.1 | 2 |
| 3 | | T0P3 | GGCTAT GGCGGC TAGGGC TAGTAT GGCTAT | 30 | 56.6 | 1 |
| 4 | | T0P4 | CTGGCA TCTGAC | 12 | 58.3 | 5 |
| 1 | TP1 | T1P1 | GCTCA GTCAGA GCCATA TGCCAG | 23 | 56.5 | 3 |
| 2 | | T1P2 | GCTCA GTCAGA CTAGCC TGCCAG | 23 | 60.8 | 2 |
| 3 | | T1P3 | GGCTAT GGCGGC TAGGGC TAGTAT GGCTAT | 30 | 56.6 | 1 |
| 4 | | T1P4 | TGAGC CTGGCA TCTGAC | 17 | 58.8 | 5 |
| 1 | TH0 | T0H1 | GTCAGA CACGAG TGCCAG | 18 | 61.1 | 2 |
| 2 | | T0H2 | GTCAGA CAGGAG TGCCAG | 18 | 61.1 | 2 |
| 3 | | T0H3 | GTCAGA CCTAGA TGCCAG | 18 | 55.5 | 2 |
| 4 | | T0H4 | GTG CTCCTG AAGTGG CTCGTG CTCCTG AAGTGG CTC | 36 | 61.1 | 1 |
| 5 | | T0H5 | CTGGCA TCTGAC | 12 | 58.3 | 6 |
| 1 | TH1 | T1H1 | TGAGCG GTCAGA CACGAG TGCCAG | 24 | 62.5 | 2 |
| 2 | | T1H2 | TGAGCG GTCAGA CAGGAG TGCCAG | 24 | 62.5 | 2 |
| 3 | | T1H3 | TGAGCG GTCAGA CCACTT TGCCAG | 24 | 58.3 | 2 |
| 4 | | T1H4 | GTG CTCCTG TCTAGG CTCGTG CTCCTG TCTAGG CTC | 36 | 61.1 | 1 |
| 5 | | T1H5 | CGCTCA CTGGCA TCTGAC | 18 | 61.1 | 6 |

*Supplementary Table 1: Sequences of various DNA supramolecular architectures used for forming rigid DNA Hydrogels.*

### ScadNano Designs: 2D

1. **DX0**

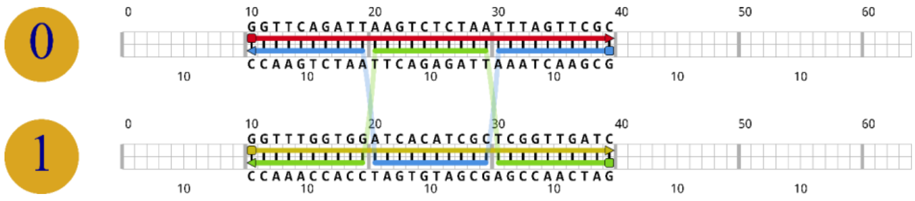

2. **DX1**

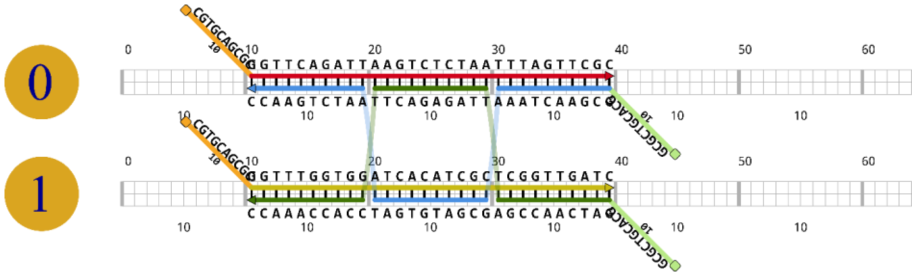

3. **DX2**

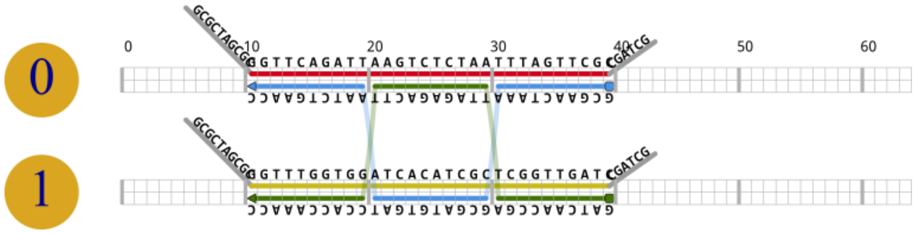

4. **PX0**

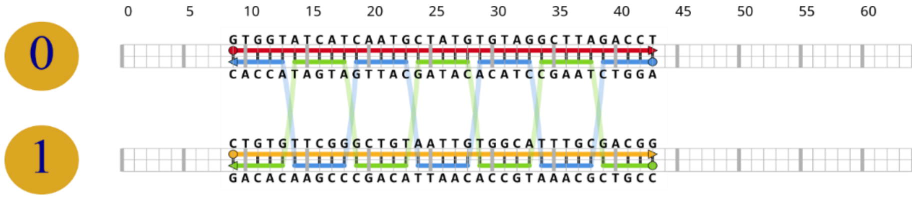

5. **PX1**

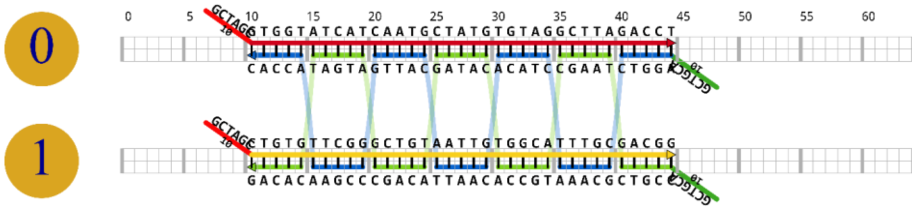

6. **PX2**

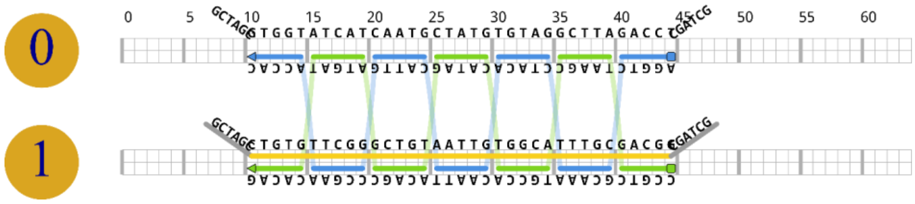

7. **TT0**

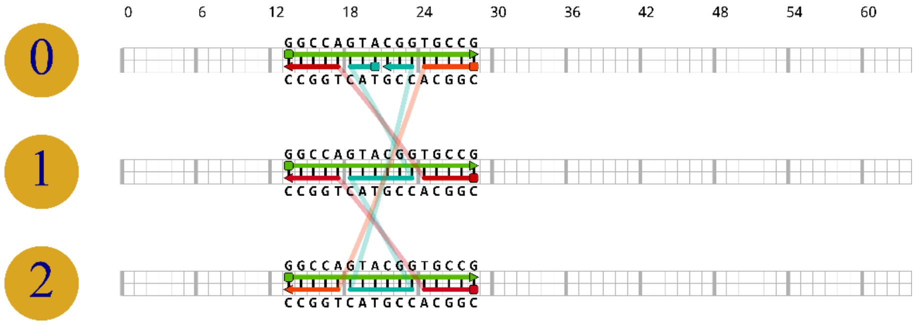

8. TT1

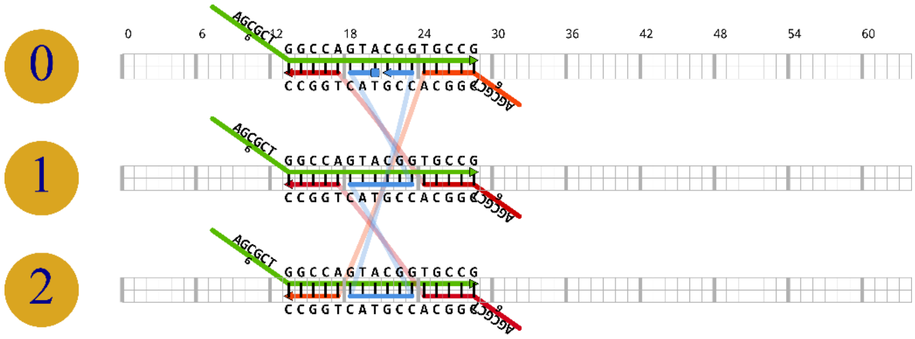

9. TQ0

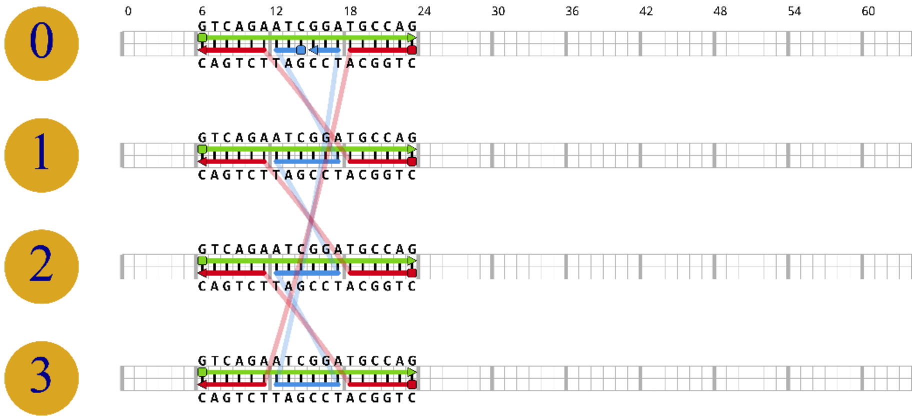

10. TQ1

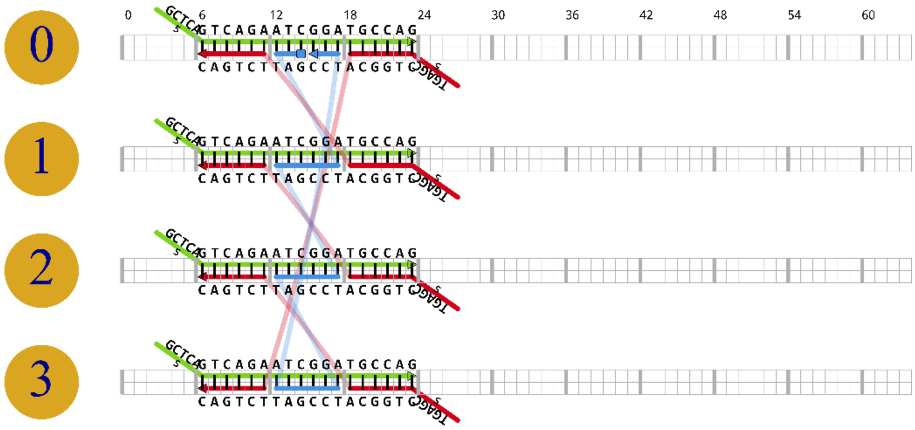

11. TP0

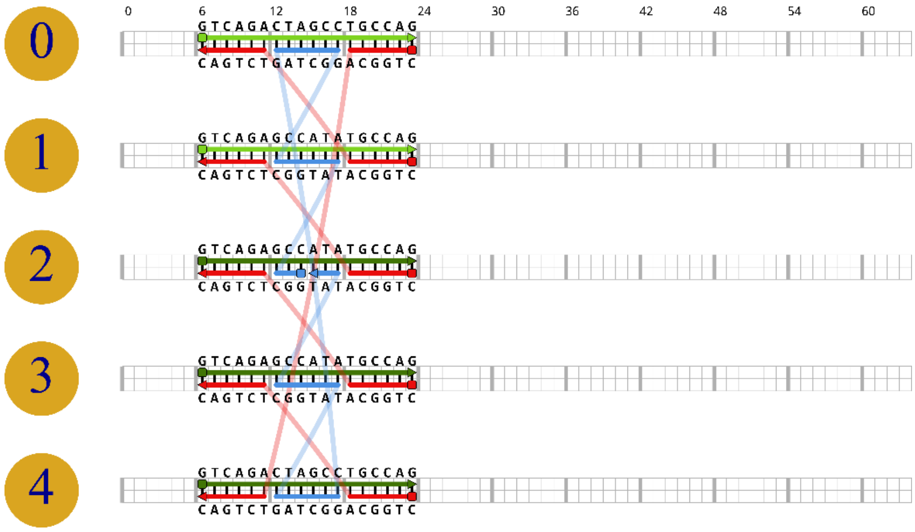

12. TP1

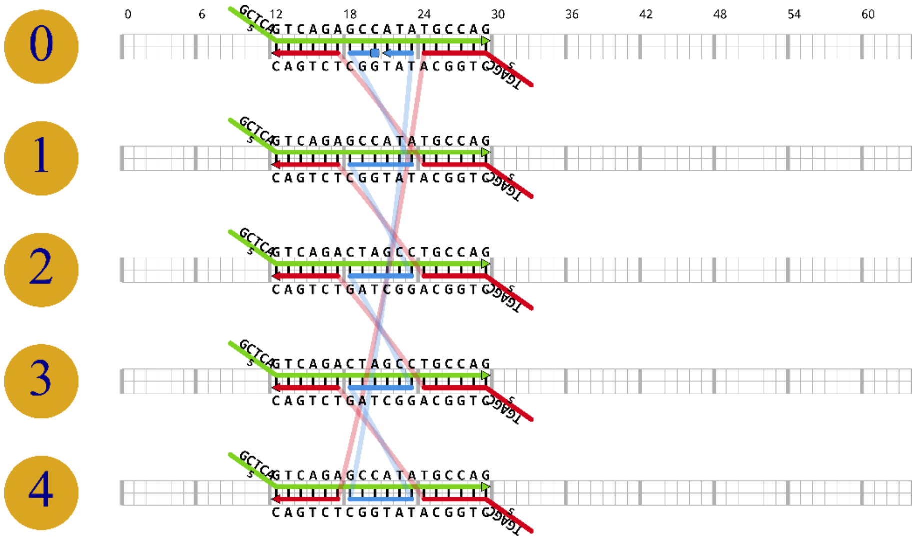

13. TH0

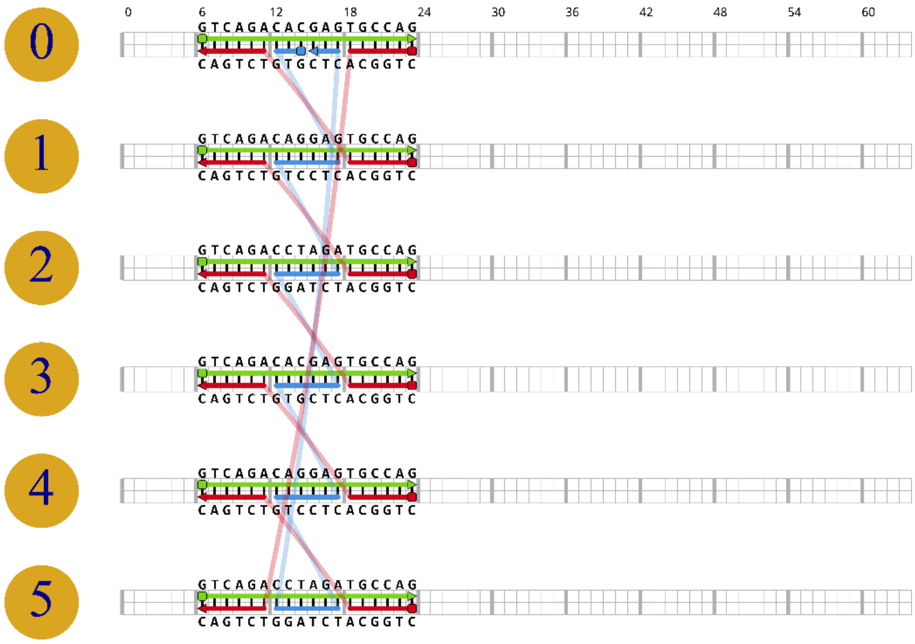

14. TH1

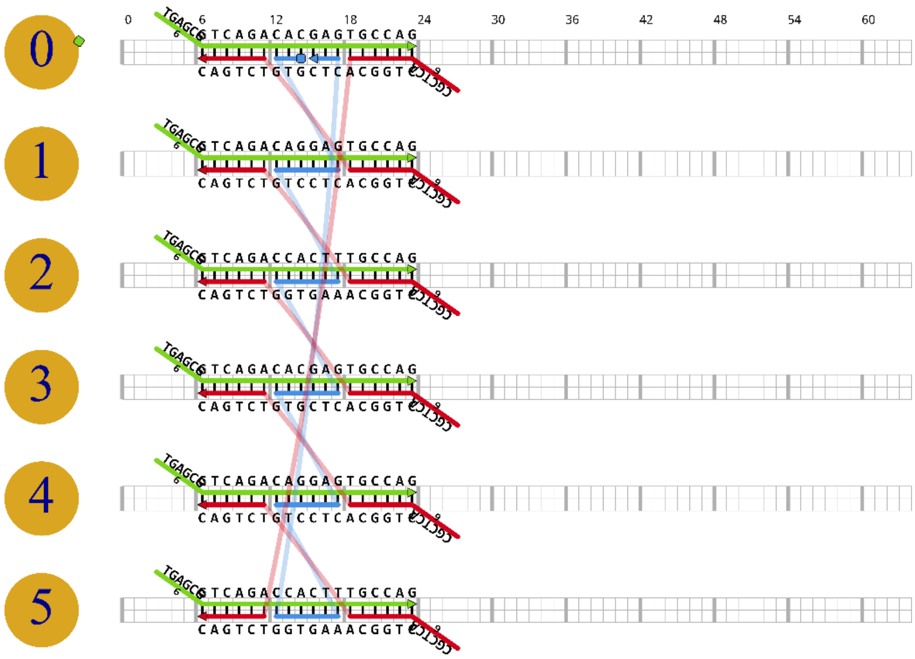

*Figure 2: 2D assembled architectures of various DNA motifs. These 2D structures were designed over the ScadNANO web server.*

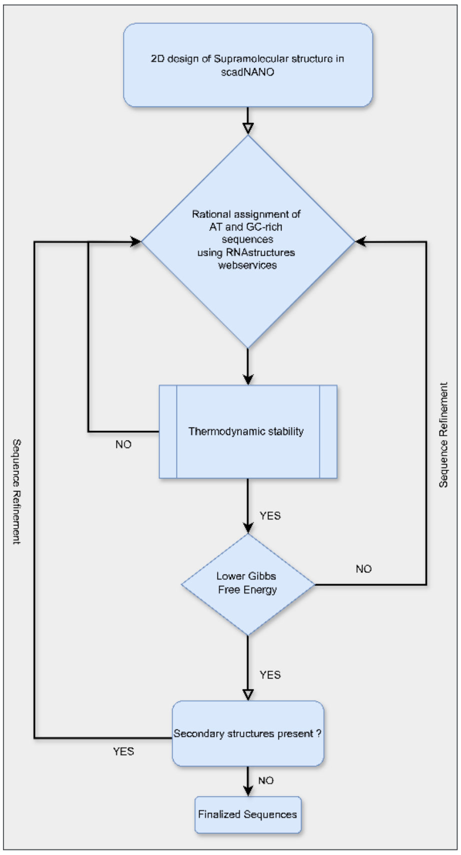

*Figure 3: A flow chart displaying the sequence design logic and pipeline.*

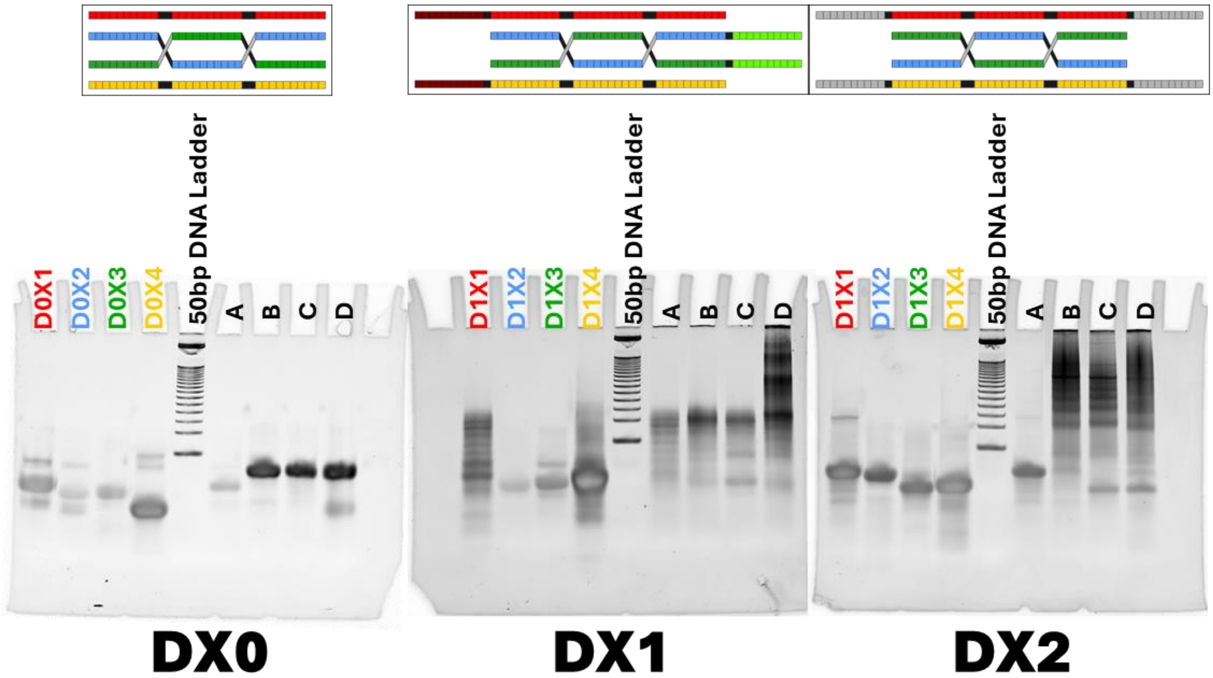

*Figure 4: EMSA band patterns of sequential assembly of ssDNA to form DX-type junction and hydrogels. The colored lanes indicate ssDNA loaded alone, while A, B, C and D indicate sequential increment of ssDNA assemblies.*

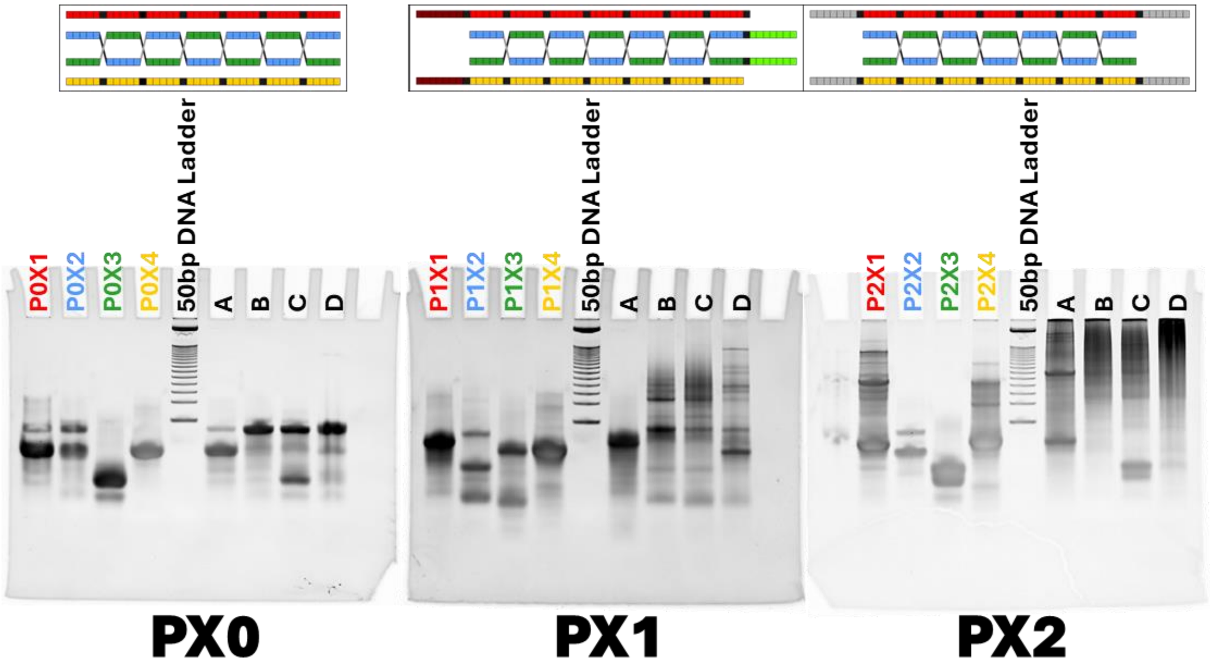

*Figure 6: EMSA band patterns of sequential assembly of ssDNA to form PX-type junctions and hydrogels. The colored lanes indicate ssDNA loaded alone, while A, B, C, and D indicate sequential increments of ssDNA assemblies.*

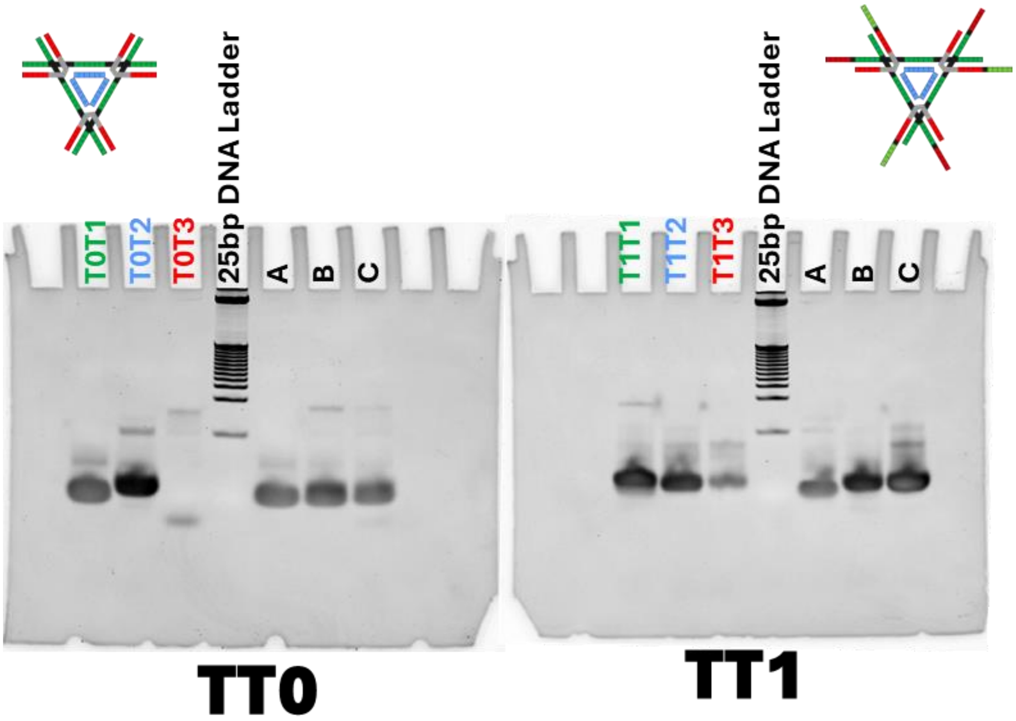

*Figure 7: EMSA band patterns of sequential assembly of ssDNA to form tensegrity triangle-type junctions and hydrogels. The colored lanes indicate ssDNA loaded alone, while A, B, and C indicate sequential increments of ssDNA assemblies.*

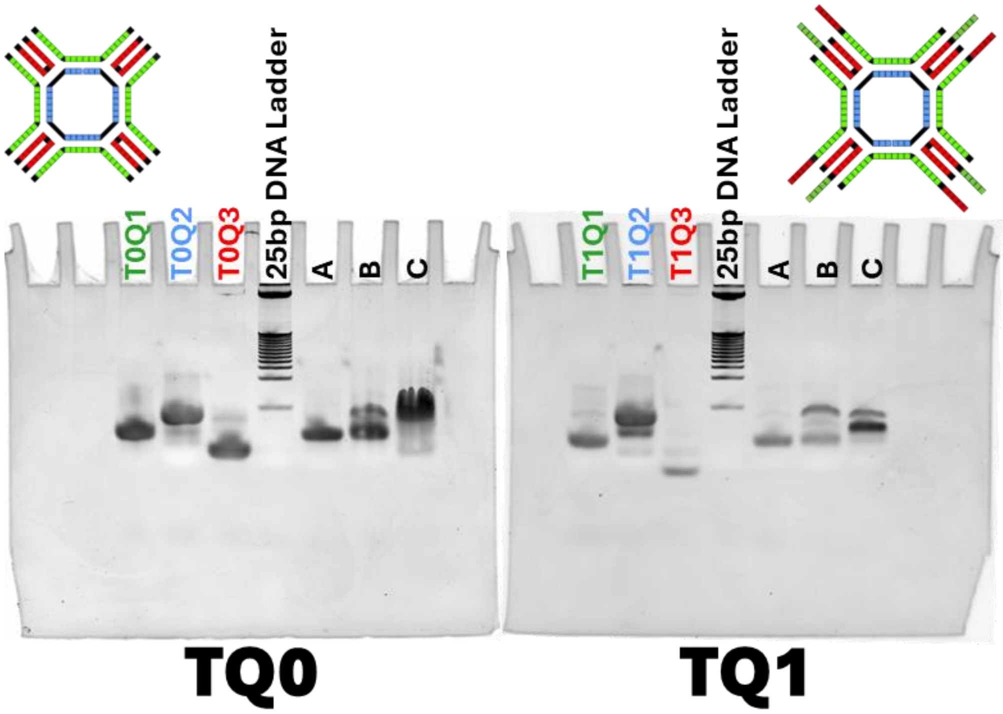

*Figure 8: EMSA band patterns of sequential assembly of ssDNA to form tensegrity Quadrilateral-type junctions and hydrogels. The colored lanes indicate ssDNA loaded alone, while A, B, and C indicate sequential increments of ssDNA assemblies.*

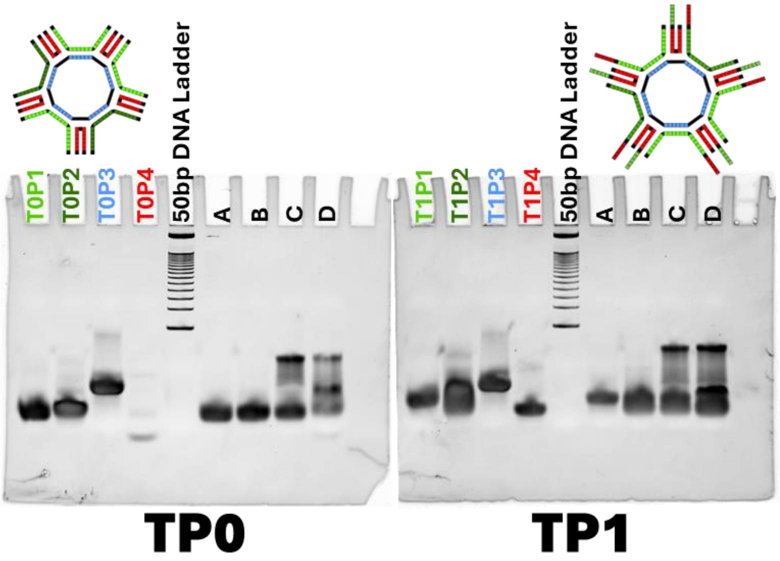

*Figure 9: EMSA band patterns of sequential assembly of ssDNA to form tensegrity Pentameric-type junctions and hydrogels. The colored lanes indicate ssDNA loaded alone, while A, B, C, and D indicate sequential increments of ssDNA assemblies.*

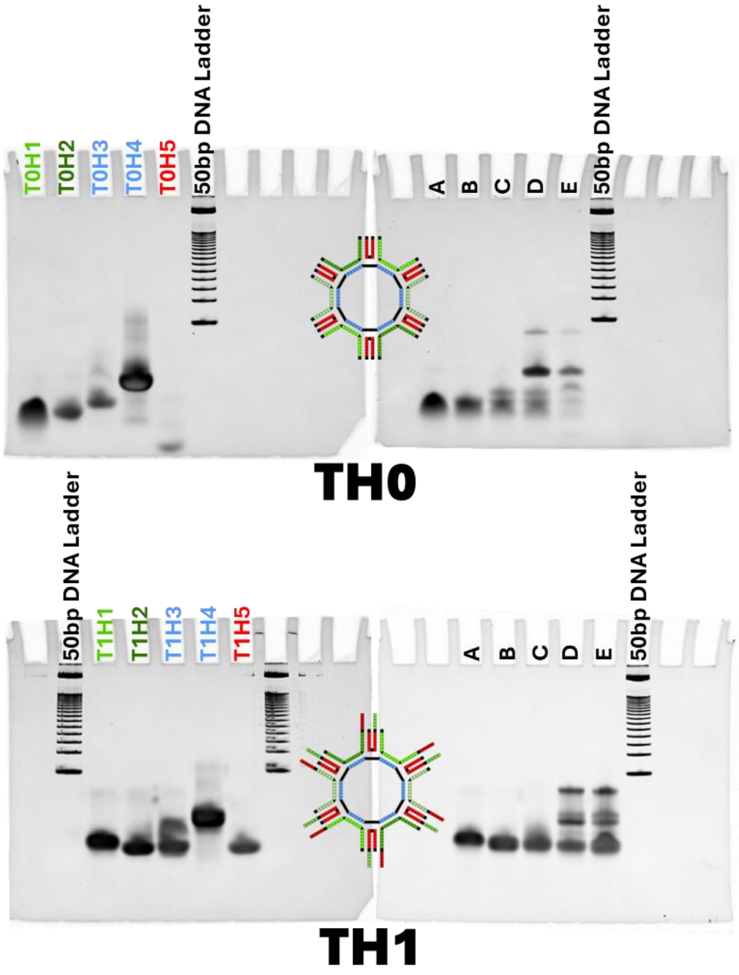

*Figure 10: EMSA band patterns of sequential assembly of ssDNA to form tensegrity Pentameric-type junctions and hydrogels. The colored lanes indicate ssDNA loaded alone, while A, B, C, and D indicate sequential increments of ssDNA assemblies.*

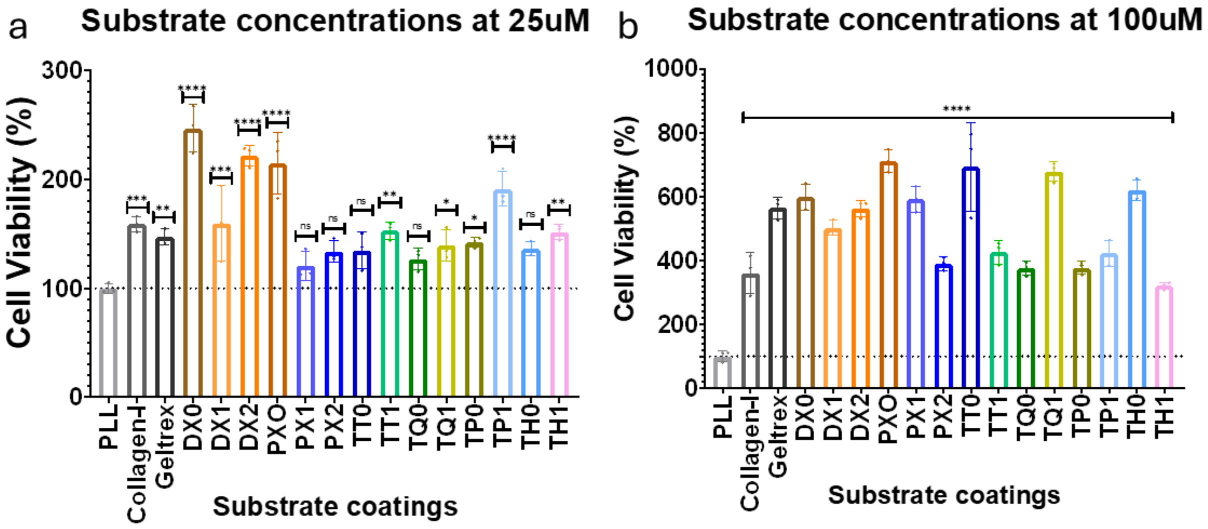

*Figure 11: Cytotoxicity assay of DNA hydrogels at concentrations of 25µM (a) and 100µM (b). The analysis was carried out by ordinary ANOVA tests. **** signifies p-value<0.0001.*

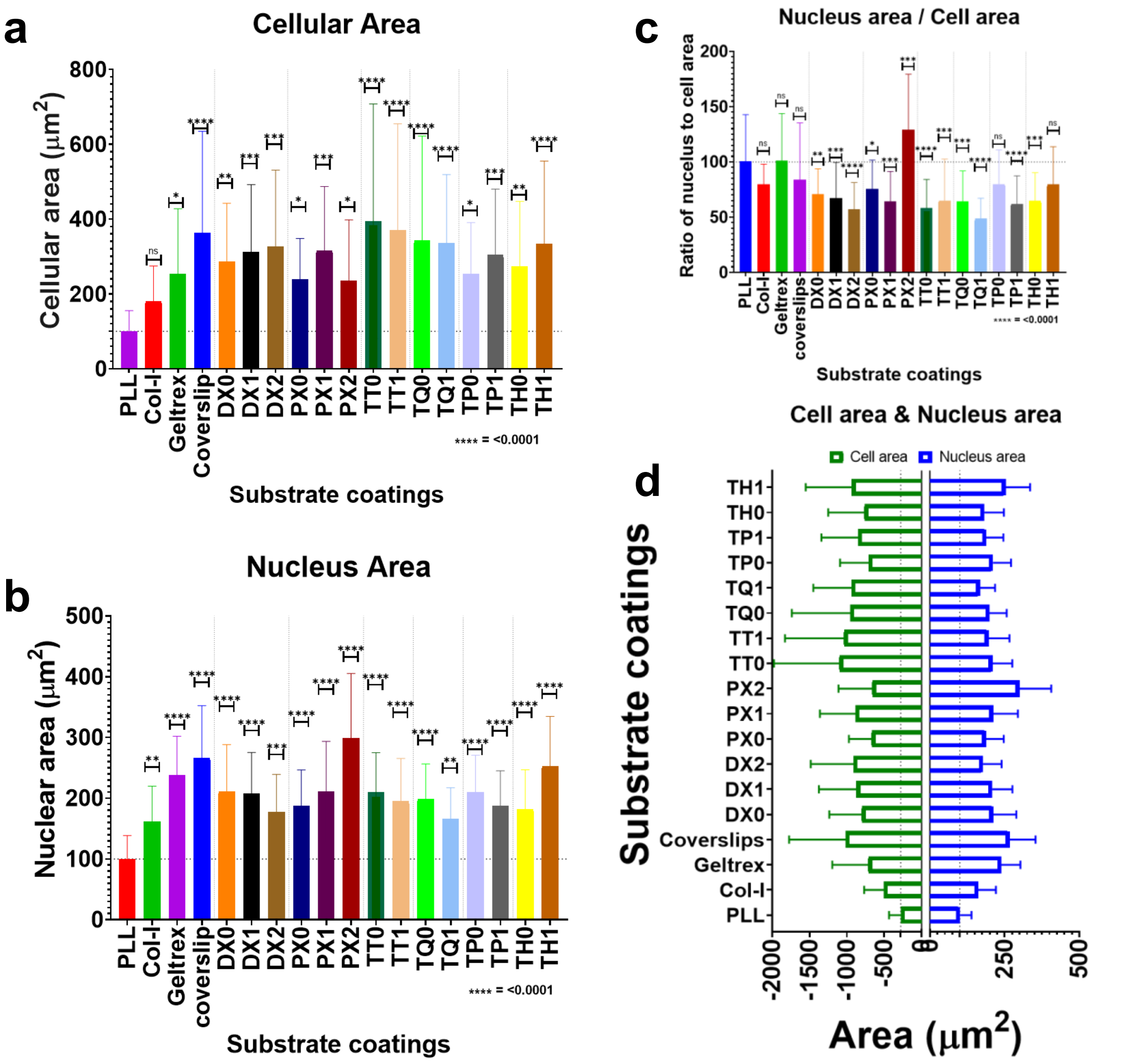

*Figure 12: Quantified data of cellular area (a) and nuclear area (b) of RPE 1 cells cultured over DNA hydrogels of 50µM concentrations. (c) Shows nuclear to cytosolic ratio, where >100 represents a cell with larger nuclei and wise verse. The analysis was carried out by ordinary ANOVA tests. **** signifies p-value<0.0001.*

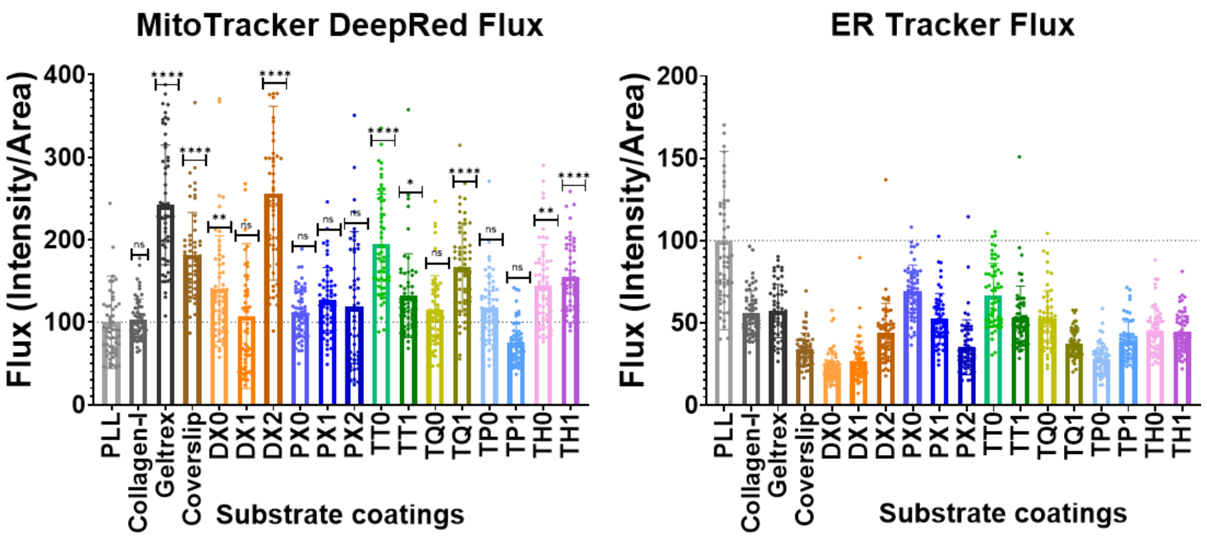

*Figure 13: Quantified data showing fluorescence intensity per unit cell area for Mitotracker DeepRed to the left and ER tracker to the right. The analysis was carried out by ordinary one-way ANOVA tests. **** signifies p-value<0.0001.*

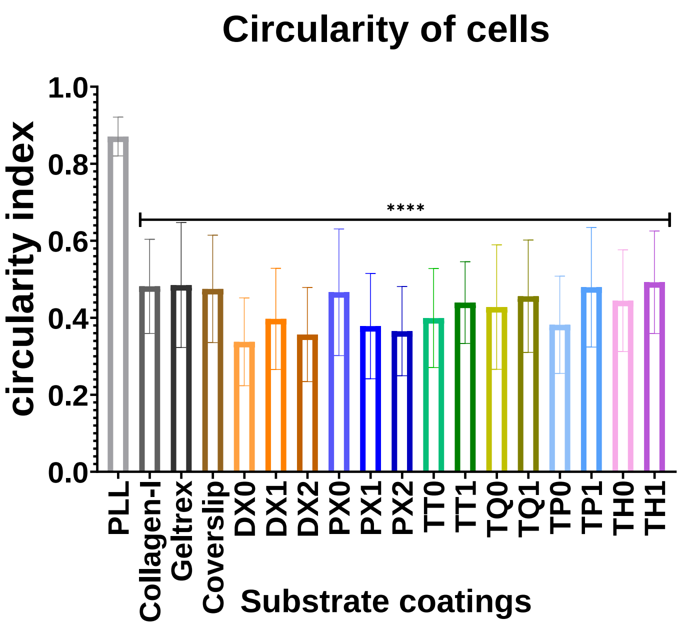

*Figure 14: The graph shows the shape descriptions of RPE 1 cells upon exposure to DNA hydrogels of varied stiffness at 50 µM concentrations. The analysis was carried out by ordinary one-way ANOVA tests. **** signifies p-value<0.0001.*
